# Selective cross-TAD gene regulation by the *Tcra* enhancer during T cell development

**DOI:** 10.64898/2026.04.27.721065

**Authors:** Alonso Rodríguez-Caparrós, Laura López-Castellanos, Jennifer López-Ros, Víctor Castro-Ruiz, María Duarte-Ruiz, Sandra Jiménez-Lozano, Adela Moreno-Castillo, Laura C. Terrón-Camero, Eduardo Andrés-León, Carlos Suñé, Cristina Hernández-Munain

**Affiliations:** Instituto de Parasitología y Biomedicina López-Neyra, Consejo Superior de Investigaciones Científicas (IPBLN-CSIC),Avenida del Conocimiento 17, 18016-Granada, Spain

**Keywords:** enhancer, transcription, Topologically Associating Domains, T-cell development, T-cell receptor

## Abstract

Long-range functional enhancer–promoter (E-P) interactions typically occur within topologically associating domains (TADs), which facilitate contacts while restricting inter-domain communication. Although approximately one-third of E-P interactions cross TAD boundaries, they are considered non-functional and are only rarely reported during embryonic development. Here, we investigated the activity of the strong *Tcra* enhancer (Eα) positioned at the boundary between a centromeric TAD containing the T-lineage-specific *Tcra-Tcrd* locus and a telomeric TAD encompassing broadly expressed *Dad1*-to-*Cdh24* genes. To directly assess Eα activity across TADs, we generated mice lacking Eα while preserving its associated CTCF-binding elements. As expected, Eα was required for T-cell specific *Tcra-Tcrd* transcription and normal T-cell development. In contrast, Eα did not activate *Dad1* or *Haus4* transcription in the telomeric TAD, indicating effective insulation by boundary elements. Unexpectedly, Eα selectively activated developmentally regulated *Cdh24* expression, the most distal gene in the telomeric TAD, in thymocytes. These findings reveal a naturally occurring, functional cross-TAD E-P interaction during adult T-cell development, demonstrating that enhancers positioned at TAD boundaries can bypass topological insulation to selectively regulate distal gene expression.

## Introduction

Enhancers are short, non-coding cis-regulatory elements that activate gene transcription in a precise spatial and temporal manner by serving as platforms for transcription factors (TFs) and facilitating the formation of enhancer-promoter (E-P) chromatin loops (Furlong & Levine, 2018). More than 100,000 enhancers are distributed throughout the genome, frequently bypassing proximal promoters to regulate genes located at considerable genomic distances (Sanyal et al, 2012; Shen et al, 2012; van Arensbergen et al, 2014). To ensure regulatory specificity, enhancers must activate their intended target genes while avoiding unintended activation of neighboring genes. Increasing evidence indicates that the genome is organized into regions of enhanced three-dimensional proximity, termed topologically associating domains (TADs) (Dixon et al, 2012; Rao et al, 2014), which arise through progressive chromatin loop extrusion (da Costa Nunes & Noordermeer, 2023; Fudenberg et al, 2016; Sanborn et al, 2015). TADs are thought to promote specific and functional E-P interactions while limiting cross-boundary communication (Phillips-Cremins et al, 2013; Rao et al, 2014; Sun et al, 2019; Symmons et al, 2014; Zhan et al, 2017), whereas TAD boundaries provide insulation, as demonstrated by targeted genomic deletions or rearrangements (Despang et al, 2019; Franke et al, 2016; Guo et al, 2015; Lupiañez et al, 2015; Narendra et al, 2015; Nora et al, 2012). These boundaries are typically defined by the recruitment of CTCF and cohesion to CTCF-binding elements (CBEs), thereby generating chromatin loops through extrusion mechanisms (de Wit et al, 2015; Dixon et al, 2012; Kim et al, 2019; Phillips-Cremins et al, 2013; Rao et al, 2014; Wutz et al, 2017; Zuin et al, 2014). Although most chromatin loops are anchored by pairs of convergently oriented CBEs, loops can also form between CBEs in the same orientation or between a CBE and regions enriched for active transcription or other chromatin-bound proteins (Busslinger et al, 2017; Jeppsson et al, 2022; Wutz et al, 2017). However, the strict insulating role of TADs have been challenged. Large-scale perturbations of TAD organization or loss of CTCF, cohesin, or associated regulators such as NIPBL or WAPL often result in relatively modest transcriptional changes (Despang et al, 2019; Ghavi-Helm et al, 2019; Nora et al, 2017; Rao et al, 2017; Schwarzer et al, 2017; Seitan et al, 2013; Zuin et al, 2014), with many E-P interactions remaining largely intact (Hsieh et al, 2022). Notably, cross-TAD E-P interactions are common, accounting for approximately one-third of all detected E-P contacts (Galupa & Crocker, 2020; Hsieh et al, 2022). Although the functional relevance of most of these interactions remains unclear, several compelling examples of functional cross-TAD regulation have been reported in recent years (Balasubramanian et al, 2024; Beccari et al, 2021; Galupa et al, 2020; Galupa et al, 2022; Hung et al, 2024; Kessler et al, 2023; Rouco et al, 2021). In particular, an elegant study in *Drosophila* embryos demonstrated functional inter-TAD E-P interactions by repositioning a *twist* enhancer to diverse genomic locations (Balasubramanian et al, 2024). In mammals, a limited number of endogenous cross-TAD regulatory events have been described during mouse embryogenesis, including regulation of limb patterning genes such as *Nell2* and *Ano6* by *Dbx2* enhancers, and control of *Pitx1* transcription by its distal Pen enhancer in E11.5-E12.5 embryos (Beccari et al, 2021; Hung et al, 2024; Rouco et al, 2021). Additional examples include long-range activation of *Hoxa2* transcription by super-enhancers in cranial neural crest cells during E10.5-E14.5 (Kessler et al, 2023), as well as altered *Xist* transcription following inversion of nearly the entire adjacent TAD in male embryonic stem cells and E1.5 female embryos (Galupa et al, 2022). Functional inter-TAD E-P communication in specific genomic contexts may arise through mechanisms such as boundary element stacking, which brings together regulatory elements located in distinct TADs (Hung et al, 2024), or through gene-rich loci that act as TAD borders and permit dynamic enhancer interactions with promoters on both sides of the boundary (Rodríguez-Carballo et al, 2017). Here, we identify a striking example of long-range transcriptional regulation mediated by an enhancer positioned at a TAD boundary, providing strong evidence for functional cross-TAD regulation in adult mammals. Together, these findings challenge the prevailing view that cross-TAD interactions are largely non-functional or, when functional, restricted to embryonic development.

The *Tcra* enhancer (Eα) is a potent T-cell-specific enhancer (Vanhille et al, 2015) that is essential for VαJα recombination and T-cell receptor (TCR) α chain (TCRα) expression (Rodríguez-Caparrós et al, 2022; Sleckman et al, 1997). Eα activity is tightly regulated during thymocyte development. It is inactive in early double-negative (DN) CD4^-^CD8^-^, specifically in the DN1 (CD25^-^CD44^+^), DN2 (CD25^+^CD44^+^), and DN3 (CD25^+^CD44^-^) subpopulations. It becomes active in DN4 (CD25^-^CD44^-^) and immature single positive CD8^+^ (ISP) thymocytes, reaches full activity in double-positive (DP) CD4^+^CD8^+^ thymocytes, and is subsequently repressed, yet remains functionally active, in single-positive (SP) CD4^+^ or CD8^+^ thymocytes and mature αβ T cells (Boudil et al, 2015; del Blanco et al, 2015; Hernández-Munain et al, 1999; Rodríguez-Caparrós et al, 2020). Eα-dependent TCRα expression in DP thymocytes permits pairing of TCRα with TCRβ to generate a functional TCRαβ complex, thereby promoting thymocyte differentiation toward the SP stage through positive selection. Structurally, Eα consists of a 116-bp enhanceosome formed by the cooperative binding of CREB/ATF2, LEF1/TCF1, Runx1, and ETS1 (Hernández-Munain et al, 1998; Hernández-Munain et al, 1999; Rodríguez-Caparrós et al, 2020; Spicuglia et al, 2000). Full enhancer activity and temporal regulation require the recruitment of additional TFs, including E2A, HEB, Ikaros, FLI1, GATA3, and RORγt, which bind within a larger ∼200-nt region (Cieslak et al, 2020; Dauphars et al, 2022; del Blanco et al, 2015; del Blanco et al, 2012; del Blanco et al, 2009; Hernández-Munain et al, 1998; Mihai et al, 2023; Naik et al, 2024; Rodríguez-Caparrós et al, 2020). Recruitment of these TFs is dynamically regulated throughout thymocyte development (Fig. S1). In DN3 thymocytes, HOXA factors interact with ETS1 bound to Eα, maintaining the enhancer in an inactive but poised state (Cieslak et al, 2020). Pre-TCR signaling in DN4 and early DP thymocytes triggers the induction of TFs such as NFAT, EGR1, EGR3, AP1, and RORγt, concomitant with dissociation of HOXA proteins, thereby enabling full enhancer activation (Cieslak et al, 2020; del Blanco et al, 2012; Naik et al, 2024). During the transition from DP to SP thymocytes, TCRαβ signaling strongly inhibits Eα activity; this repression is maintained in mature αβ T lymphocytes and correlates with reduced recruitment of E2A and HEB, as well as the absence of DP-specific RORγt (del Blanco et al, 2015; Mihai et al, 2023; Naik et al, 2024; Rodríguez-Caparrós et al, 2022). Despite this inhibition, Eα remains required for transcription and expression of the rearranged TCRα gene in both mouse and human mature T cells (Rodríguez-Caparrós et al, 2022; Schönberg et al, 2025; Sleckman et al, 1997).

Eα is sited at a conserved boundary between two TADs (Fig. 1). The ∼425-kb centromeric TAD containa T-cell specific *Tcra*-*Tcrd* gene segments extending up to *Trav17*, whereas the ∼420-kb telomeric TAD harbors unrelated, broadly expressed genes ranging from *Dad1* to *Cdh24* genes (Vanhille et al, 2015). Previous studies of Eα function relied on a mouse deletion encompassing both Eα and its two telomeric flanking CTCF binding elements (EαCBEs), Eα+EαCBE^-/-^ (Rodríguez-Caparrós et al, 2022; Sleckman et al, 1997; Zhao et al, 2020). These studies demonstrated that the combined Eα+EαCBEs region is essential for *Tcra-Tcrd* transcription, *Trav*-*Traj* (Vα-Jα) recombination, TCRα and TCRαβ expression, and αβ T-cell development. Notably, this deletion also resulted in reduced transcription of several genes across the telomeric TAD during T-cell development, including *Dad1, Haus4*, and *Cdh24*. More recently, comparison of mice lacking Eα+EαCBE with mice carrying a deletion of only the two EαCBEs, EαCBE^-/-^, led to the conclusion that EαCBEs insulate proximal telomeric genes, such as *Dad1*, from the activating influence of Eα, while being required for high expression of more distally located genes, such as *Cdh24* (Zhao et al, 2020). Here, we directly investigated the role of Eα in both TADs by generating a mouse allele in which the enhancer is deleted while its flanking EαCBEs are preserved (Eα ^-/-^). As expected, deletion of Eα alone recapitulated the well-established effects of the Eα+CBE deletion on *Tcra-Tcrd* transcription and T-cell development. Unexpectedly, however, although loss of Eα did not substantially affect the expression of most genes within the telomeric TAD when EαCBEs remained intact, transcription of the distal *Cdh24* gene was found to be highly dependent on Eα. This study provides a striking example of enhancer-mediated regulation of distal genes across TAD boundaries beyond embryogenesis (Beccari et al, 2021; Hung et al, 2024; Kessler et al, 2023; Rouco et al, 2021), offering strong evidence for long-range transcriptional regulation expression by an enhancer positioned at a TAD border. These findings support the hypothesis that enhancers located at domain boundaries can exploit this position to contact and activate genes in neighboring TADs (Hung et al, 2024), challenging the prevailing view that TADs rigidly confine enhancer activity within individual domains and underscoring the broader regulatory potential of boundary-positioned enhancers throughout mammalian development.

**Figure 1.**
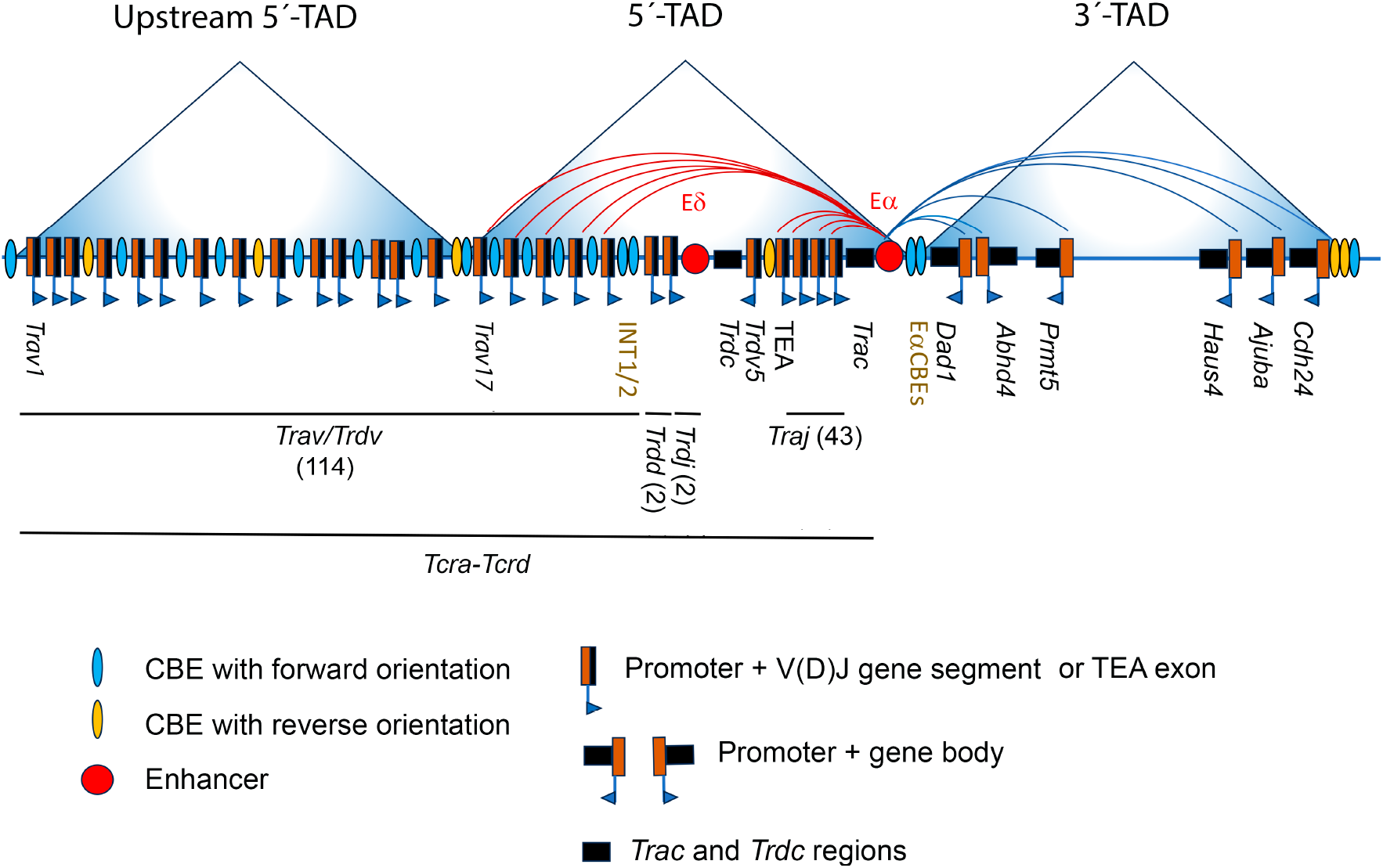
Schematic representation of *Tcra* E-P interactions across flanking TADs in DP thymocytes. The diagram illustrates transcriptionally active E-P interactions mediated by Eα within the centromeric TAD (red arcs), and the telomeric TAD (blue arcs). The *Tcra*-*Tcrd* locus comprises variable (*Trav* and *Trdv*), diversity (*Trdd*), and joining (*Traj* and *Trdj*) gene segments, depicted as black rectangles, each preceded by its corresponding promoter (brown rectangles). The number of functional segments is indicated. Constant regions are also shown as black rectangles. The T early α (TEA) exon and its promoter are labeled. The enhancers Eα and Eδ are represented as red circles. Transcript orientation is indicated by blue arrows at promoter sites. CTCF binding elements (CBEs) are depicted as blue or orange ovals to denote orientation. Genes located within the telomeric TAD and relevant to this study are also annotated.

## Results

### Generation of two independent Eα^-/-^ mouse lines

The 1.1-kb Eα+EαCBE region is known to be essential for *Tcra-Tcrd* transcription and *Tcra* recombination within the centromeric TAD, but it also influences gene transcription within the telomeric TAD genes (Rodríguez-Caparrós et al, 2022; Sleckman et al, 1997; Zhao et al, 2020). In particular, deletion of this region results in a marked reduction of *Dad1* and *Haus4* transcript levels in DN3 and DP thymocytes, as well as in non-T tissues such as liver, kidney, and testis, and a decrease of *Cdh24* transcription in DP thymocytes (Rodríguez-Caparrós et al, 2022; Zhao et al, 2020). To determine whether the transcriptional effects observed in Eα+EαCBE^-/-^ mice on telomeric TAD genes were specifically attributable to loss of the enhancer itself, we generated two independent Eα-deficient mouse lines carrying a precise 200-bp deletion of the Eα region. These lines, designated Eα1^-/-^ and Eα2^-/-^, harbor similar but not identical deletions (Figs. S2 and S3) and were backcrossed onto the wild type (WT) C57bl/6 background to minimize potential off-target effects and ensure reproducibility. Both lines exhibited indistinguishable phenotypes across all assays performed (Figs. S4, S5, S8 and S9). Data presented in the main figures were obtained using Eα1^-/-^ mice, hereafter referred as Eα ^-/-^, whereas results from both mouse lines are included in the Supplementary Information.

### Indistinguishable T-cell developmental phenotypes in Eα+EαCBE^-/-^ and Eα^-/-^ mice

αβ T cell maturation is severely impaired in Eα+EαCBE^-/-^ mice as a result of a developmental block at the DP thymocyte stage, despite normal thymic cellularity, due to defective positive selection (Rodríguez-Caparrós et al, 2022; Sleckman et al, 1997). Comparative flow-cytometric analyses revealed that thymocyte development in Eα+EαCBE^-/-^ and Eα^-/-^ thymocytes was indistinguishable, with similar proportions of DN, DP, and SP thymocytes, as well as comparable CD3 expression patterns (Figs. 2 and S4). Both mutant strains displayed a pronounced arrest at the DP stage, characterized by markedly reduced CD3 (Figs. 2A, 2B, S4A, and S4B). This effect results from failure to express TCRα and, consequently, the TCRαβ heterodimer, thereby preventing positive selection of TCRαβ-expressing DP thymocytes. In agreement with these findings, analysis of splenocyte populations did not reveal any differences between Eα+EαCBE^-/-^ and Eα^-/-^ mice (Figs. 2C-2E and S4C-S4E). In both mutant models, splenic analyses revealed a dramatic reduction in αβ T cells, as evidenced by a significant decrease in the CD3^+^ population. Concomitantly, both Eα^-/-^ and Eα+EαCBE^-/-^ mice exhibited an increased proportion of γδ T cells, accompanied by a corresponding reduction in αβ T cells. Consistent with the predominance of Vα2^+^ cells among αβ T cells in Eα+EαCBE^-/-^ mice, arising from Eδ-dependent *Trav14* rearrangements (Aifantis et al, 2006; Rodríguez-Caparrós et al, 2022; Sleckman et al, 1997), a comparable distribution of Vα2^+^ cells was also observed in Eα^-/-^ mice. In line with the requirement for Eα in achieving high-level expression of rearranged *Tcra* and *Tcrd* in mature T cells (Rodríguez-Caparrós et al, 2022; Sleckman et al, 1997), surface expression of Vα2 and TCRγδ was similarly reduced in splenocytes from both Eα+EαCBE^-/-^ and Eα^-/-^ mice relative to WT controls (Figs. 2E and S4E). This reduction indicates a decreased number of TCR complexes expressed at the cell surface of both αβ and γδ T lymphocytes. Collectively, these results demonstrate that Eα+EαCBE^-/-^ and Eα^-/-^ mice exhibit indistinguishable defects in T-cell development, confirming that the genomic region deleted in Eα^-/-^ mice encompasses all the sequences required for proper TCRα and TCRδ expression during thymocyte differentiation.

**Figure 2.**
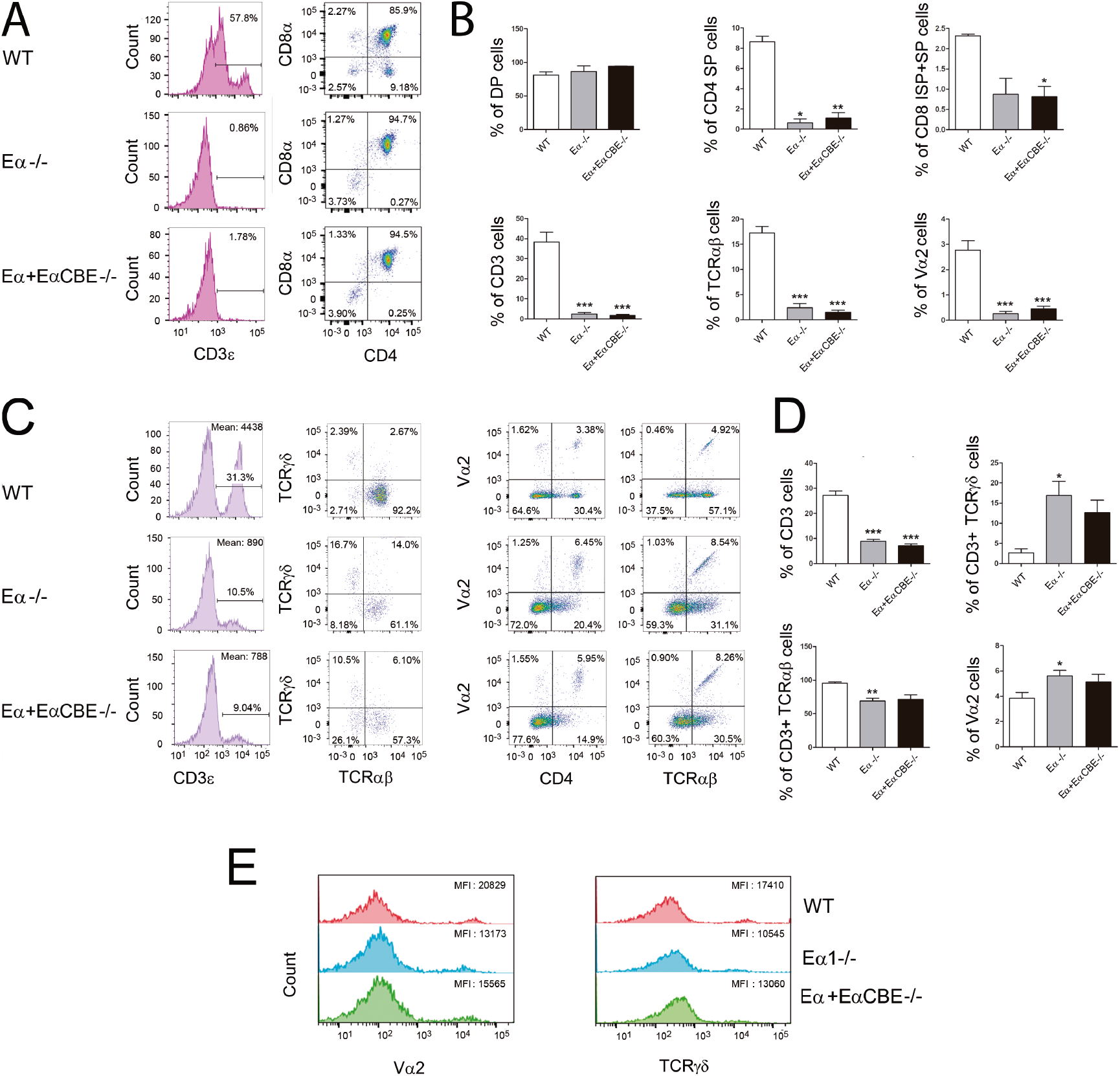
Indistinguishable T-cell development phenotypes in Eα^-/-^and Eα+EαCBE^-/-^ mice. **(A)** Representative flow cytometry plots of total thymocytes from the indicated genotypes are shown, with percentages of cells in the relevant quadrants indicated. **(B)** Percentages of the indicated thymocyte populations are presented as mean ± SEM from 3-10 independent experiments, using 4-5 mice per genotype. **(C)** Representative flow cytometry plots of total splenocytes from the indicated genotypes are shown, with percentages of cells in the relevant quadrants indicated. Right panels present TCRαβ and TCRγδ expression among CD3^+^ thymocytes shown in the respective left panels. **(D)** Percentages of the indicated splenocyte populations are shown as mean ± SEM from 3-10 independent experiments, using 4-5 mice per genotype. **(E)** Representative flow cytometry plots showing Vα2 and TCRγδ expression in splenocytes from the indicated mouse strains are shown; mean fluorescence intensities (MFIs) are indicated. Statistical analyses were performed using a non-parametric unpaired Student’s *t* test with Welch correction comparing mutant data with WT controls: \**p* < 0.05, \*\**p* < 0.005, and \*\*\**p* < 0.0005.

### Both Eα and Eα+EαCBE deletions inhibit Tcra-Tcrd germline transcription in DP thymocytes and rearranged Tcrd transcription in γδ T cells

Deletion of Eα+EαCBE region has been shown to be essential for transcription of both unrearranged (germline) and rearranged *Tcra* alleles during T-cell development (Rodríguez-Caparrós et al, 2022; Sleckman et al, 1997). To directly compare the impact of deleting the enhancer alone versus deleting the enhancer together with its flanking EαCBEs, we first assessed global *Tcra* gene transcription by quantifying total *Trac* (Cα) transcripts in total WT, Eα+EαCBE^-/-^, and Eα^-/-^ thymocytes (Figs. 3A and S5A). In all three genotypes, thymocyte populations are predominantly composed of DP thymocytes (Figs. 2A and S4A). Notably, Cα transcripts were completely undetectable in both mutant strains, indicating that Eα itself is required for *Tcra* transcription. Because *Tcra* undergoes Vα-Jα recombination in WT DP thymocytes, but remains largely unrearranged in Eα+CBE-deleted mice (Rodríguez-Caparrós et al, 2022), the absence of Cα transcripts in Eα+EαCBE^-/-^ and Eα^-/-^ thymocytes strongly suggested that Eα is required for transcription of the unrearranged *Tcra* locus. To formally test this hypothesis, we analyzed spliced Jα61-Cα, Jα58-Cα, and total Cα transcripts in V(D)J-recombination-deficient thymocytes from PBS- or anti-CD3ε-treated *Rag2*^-/-^, *Rag2*^-/-^ x Eα+EαCBE^-/-^, and *Rag2*^-/-^ x Eα^-/-^ mice (Fig. 3B). Injection of anti-CD3ε antibody efficiently induces DN3-to-DP differentiation in *Rag2*^-/-^ mice within 7-10 days (Rodríguez-Caparrós et al, 2019; Shinkai & Alt, 1994). Accordingly, *Rag2*^-/-^ thymi consisted almost exclusively of DN3 thymocytes (∼99.4% of total thymocytes), whereas anti-CD3ε-treated *Rag2*^-/-^ thymi contained predominantly DP thymocytes (∼95.0% of total thymocytes). As expected, DN3 thymocytes from PBS-injected *Rag2*^-/-^ mice lacked detectable *Tcra* germline transcription, whereas DP thymocytes from antibody-injected *Rag2*^-/-^ mice displayed abundant *Traj61*-Cα, *Traj58*-Cα, and total Cα transcripts. In contrast, all these transcripts were undetectable in DP thymocytes from both anti-CD3ε-treated *Rag2*^-/-^ mice carrying either the Eα+CBE or Eα deletion, demonstrating that Eα contains the essential cis-regulatory elements required for germline *Tcra* transcription.

**Figure 3.**
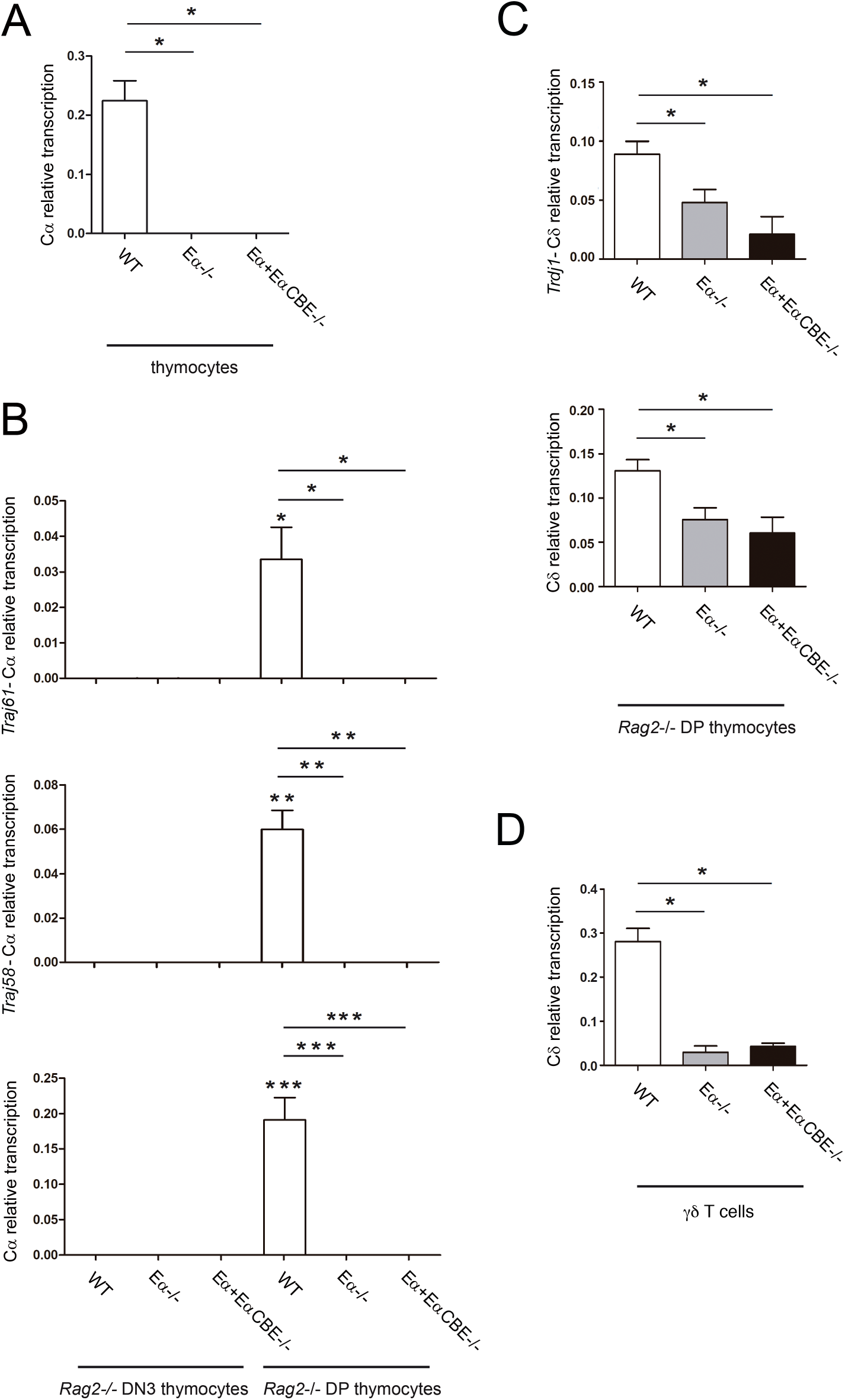
Analysis of Eα-dependent *Tcra-Tcrd* transcription. **(A)** RT-qPCR analysis of Cα transcripts in total thymocytes from WT, Eα^-/-^, and Eα+EαCBE^-/-^ mice. **(B)** RT-qPCR analysis of the indicated *Tcra* transcripts in total thymocytes from and PBS- or anti-CD3ε-injected *Rag2*^-/-^, *Rag2*^-/-^ x Eα^-/-^, and *Rag2*^-/-^ x Eα+EαCBE^-/-^ mice. **(C)** RT-qPCR analysis of the indicated *Tcrd* transcripts in anti-CD3ε-injected *Rag2*^-/-^, *Rag2*^-/-^ x Eα ^-/-^, and *Rag2*^-/-^ x Eα+EαCBE^-/-^ mice. **(D)** RT-qPCR analysis of Cδ transcripts in γδ T cells isolated from the spleens of WT, Eα ^-/-^, and Eα+EαCBE^-/-^ mice. Gene expression levels were normalized to *Actb* and are presented as mean ± SEM of duplicate measurements from 3-8 independent experiments. Statistical comparisons were performed using a non-parametric unpaired Student’s *t* test with Welch correction, as indicated: \**p* < 0.05, \*\**p* < 0.005, and \*\*\**p* < 0.0005.

Previous studies have also established a critical role for Eα in regulating unrearranged *Tcrd* transcription in DP thymocytes, as well as normal transcription of rearranged *Tcrd* alleles in γδ T cells (Hernández-Munain et al, 1999; Sleckman et al, 1997). Analysis of total *Trdc* (Cδ) transcript levels in thymocytes from WT, Eα+EαCBE^-/-^, and Eα^-/-^ mice revealed a marked inhibition of *Tcrd* transcription in both mutant strains (Fig. S5B). However, because *Tcrd* is deleted from the chromosome during αβ T-cell development in the vast majority of WT thymocytes, it cannot be regulated by Eα on most alleles in these cells. Moreover, measurement of Cδ transcripts in total thymocytes does not distinguish between unrearranged (germline) and rearranged *Tcrd* transcripts. To specifically assess germline *Tcrd* transcription, we therefore analyzed DN3 and DP thymocytes derived from PBS- or anti-CD3ε-treated *Rag2*^-/-^ mice carrying either the Eα or Eα+EαCBE deletion (Figs. 3C and S5C). Both deletions resulted in a comparable and pronounced inhibition of germline *Trdj1*-Cδ and Cδ transcripts in DP thymocytes (Fig. 3C), whereas no effect of these deletions was observed in DN3 thymocytes (Fig. S5C). Furthermore, analysis of rearranged *Tcrd* transcription in purified splenic γδ T cells revealed a similarly strong reduction of Cδ transcripts in both mutant genotypes (Fig. 3D).

Together, these results clearly demonstrate that deletion of either Eα alone or the Eα+EαCBE region results in equivalent inhibition of both germline *Tcra* and *Tcrd* transcription in DP thymocytes, as well as rearranged *Tcrd* transcription in γδ T cells, establishing that the Tα1-Tα4 region is essential for proper regulation of the *Tcra-Tcrd* locus during T-cell development.

### Eα does not activate *Dad1* and *Haus4* transcription

In addition to its role in germline *Tcra-Tcrd* transcription, deletion of the Eα+EαCBE^-/-^ region has been reported to cause a strong reduction in *Dad1* and *Haus4* transcription within the telomeric TAD in both thymocytes and non-T cells (Rodríguez-Caparrós et al, 2022; Sleckman et al, 1997; Zhao et al, 2020). Dad1 is involved in N-linked glycosylation and protects cells from apoptotic cell death, whereas Haus4 is required for mitotic spindle assembly and maintenance of centrosome integrity during cell division (Kelleger & Gilmore, 1997; Lawo et al, 2009). Given that both genes are essential for early embryonic development (Brewster et al, 2000; Hong et al, 2000) (https://www.mousephenotype.org/), it was unexpected that Eα+EαCBE^-/-^ mice exhibit no overt developmental abnormalities beyond the severe defect in αβ T-cell development (Rodríguez-Caparrós et al, 2022; Sleckman et al, 1997). To determine whether reduced transcript levels of *Dad1* and *Haus4* in Eα+EαCBE^-/-^ mice translate into decreased protein expression, we performed Western blot analyses of DN3 thymocytes from *Rag2*^-/-^ and *Rag2*^-/-^ x Eα+EαCBE^-/-^ mice, DP thymocytes from *Rag2*^-/-^ x TCRβtg and *Rag2*^-/-^ x TCRβtg x Eα+EαCBE^-/-^ mice, as well as liver, kidney, and testis tissues from *Rag2*^-/-^ and *Rag2*^-/-^ x Eα+EαCBE^-/-^ mice (Fig. S6). In all tissues analyzed, Dad1 and Haus4 protein levels were comparable between genotypes, indicating that post-transcriptional mechanisms likely compensate for reduced mRNA levels and prevent deleterious consequences of transcriptional downregulation.

To specifically evaluate the contribution of Eα itself to transcriptional regulation within the telomeric TAD, we first confirmed by ChIP-qPCR that deletion of Eα does not affect CTCF recruitment to the flanking EαCBEs in thymocytes (Figs. 4 and S7). Using primers that amplify a region preserved in all three genotypes and located downstream of the EαCBEs, we detected comparable levels of CTCF occupancy at EαCBEs in WT and Eα^-/-^ thymocytes (Fig. 4). This result was independently confirmed using primers mapping directly to the EαCBE region, which revealed similar levels of CTCF recruitment in thymocytes from WT and Eα^-/-^ strains (Fig. S7). We then compared *Dad1* and *Haus4* transcript levels in thymus and non-T tissues (kidney, liver, and testis) from WT, Eα+EαCBE^-/-^, and Eα^-/-^ mice (Figs. 5A, 5B, S8A and S8B). While transcription was markedly reduced in Eα+EαCBE^-/-^ mice, Eα^-/-^ mice showed transcript levels comparable to those in WT controls in all tissues. Further analysis of DN3 and DP thymocytes from PBS- and CD3ε-antibody-injected *Rag2*^-/-^, *Rag2*^-/-^ x Eα+EαCBE^-/-^, and *Rag2*^-/-^ x Eα^-/-^ mice, respectively, confirmed these findings (Fig. 5C). These results are consistent with the distinct regulatory mechanisms governing *Dad1* and *Haus4* compared to Eα-dependent *Tcra* transcription during thymocyte development (Fig. S8C). Together, these results demonstrate that deletion of Eα alone, in the presence of intact EαCBEs, does not disrupt transcription of the telomeric TAD genes *Dad1* and *Haus4*.

**Figure 4.**
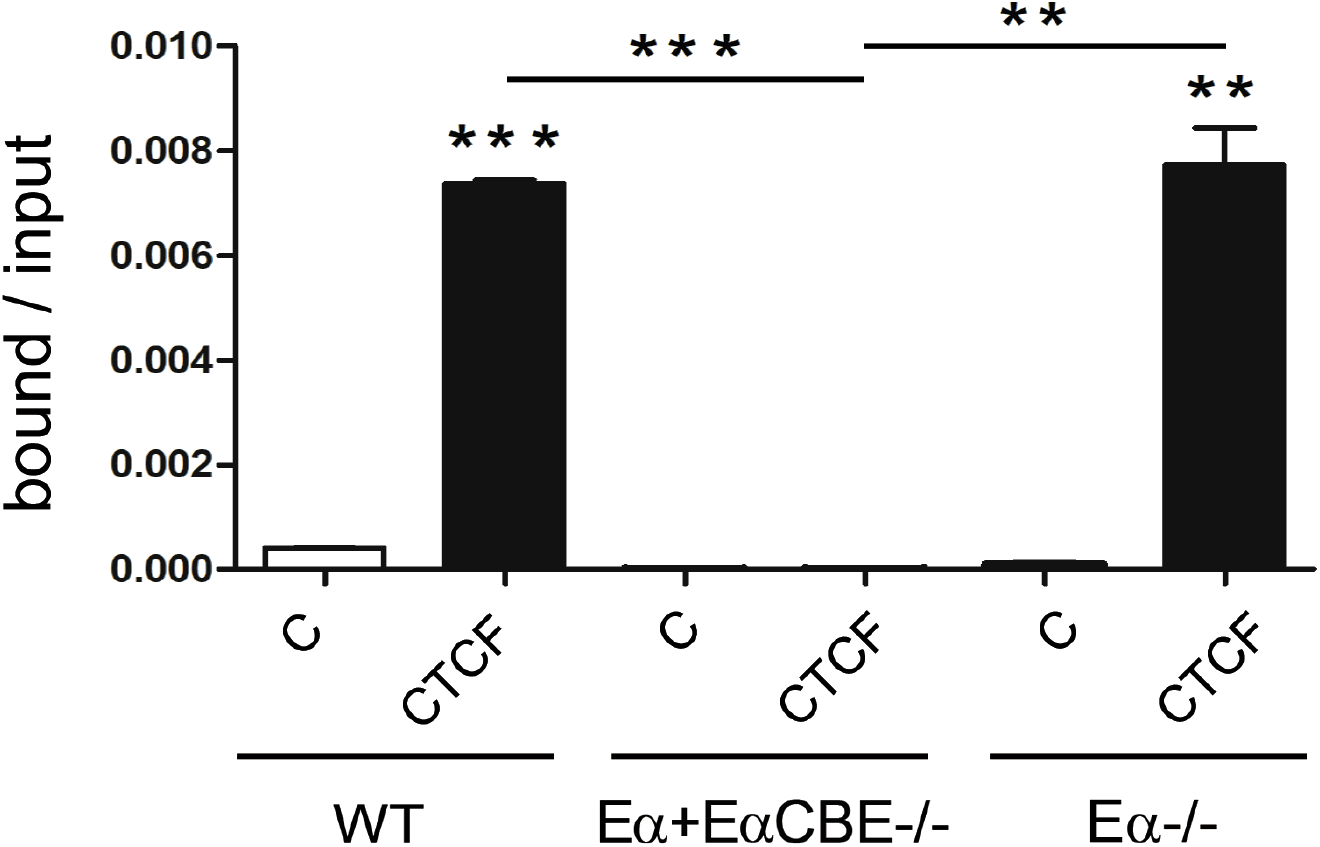
Eα^-/-^ mice retain intact EαCBEs in thymocytes. CTCF binding to the EαCBEs was assessed by ChIP-qPCR in thymocytes from WT, Eα^-/-^, and Eα+EαCBE^-/-^ mice using primers that amplify a region located downstream of the EαCBE region and preserved in all three genotypes. Data are presented as mean ± SEM from three independent experiments. Statistical analyses were performed using a non-parametric unpaired Student’s *t* test with Welch correction: \**p* < 0.05, and \*\**p* < 0.005. Statistical significance between samples immunoprecipitated with control antibody (C) and anti-CTCF antibody (CTCF) is indicated.

**Figure 5.**
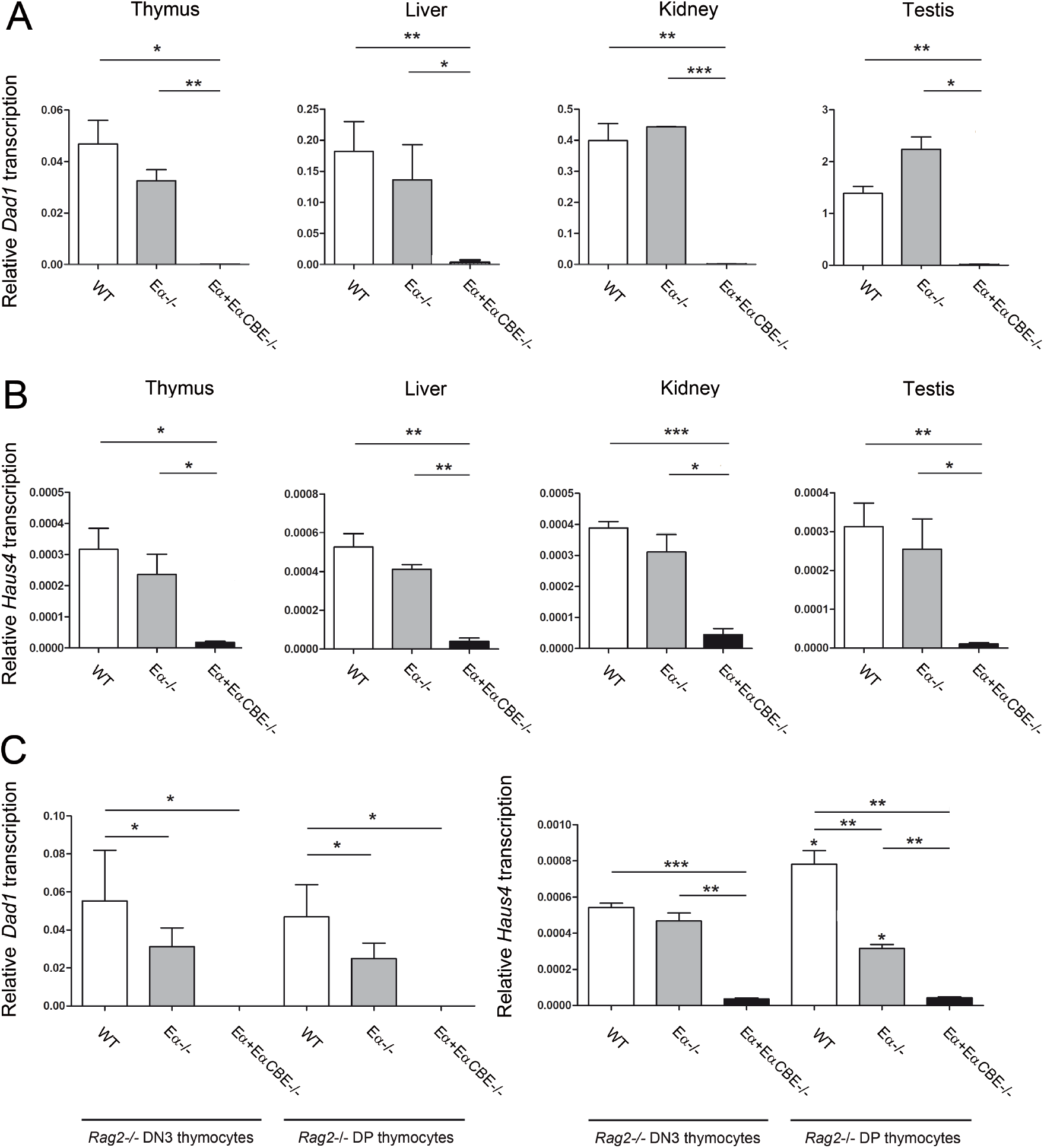
Eα-independent transcription of *Dad1* and *Haus4*. **(A)** RT-qPCR analysis of *Dad1* transcripts in thymus, liver, kidney, and testis from WT, Eα+EαCBE^-/-^, and Eα^-/-^ mice (n=3-4). **(B)** RT-qPCR analysis of *Haus4* transcripts in thymus, liver, kidney, and testis from WT, Eα+EαCBE^-/-^, and Eα^-/-^ mice (n=3-4). **(C)** RT-qPCR analysis of *Dad1* and *Haus4* transcription in PBS-injected and anti-CD3ε-injected *Rag2*^-/-^, *Rag2*^-/-^ x Eα+EαCBE^-/-^, and *Rag2*^-/-^ x Eα^-/-^ mice, as indicated (n=4-6). Gene expression levels were normalized to *Actb* and presented as mean ± SEM of duplicate measurements from the indicated number of independent experiments. Statistical comparisons were performed using a non-parametric unpaired Student’ *t* test with Welch correction, as indicated. Significance levels are indicated as follows: \**p* < 0.05, \*\**p* < 0.005, and \*\*\**p* < 0.0005.

### Eα drives *Cdh24* transcription despite the presence of EαCBEs

Unlike *Dad1* and *Haus4*, transcription of the most distal telomeric TAD gene *Cdh24* was similarly reduced in both Eα+EαCBE^-/-^ and Eα^-/-^ thymocytes (Figs. 6A and S9A). Notably, RNA-seq data indicate that *Cdh24* transcription increases during thymocyte development, reaching maximal levels in DP thymocytes and becoming strongly repressed in mature T cells (Fig. 6B) (www.immgen.org). This developmental expression pattern closely mirrors that of Eα-dependent germline *Tcra* transcription (del Blanco et al, 2015; Hernández-Munain et al, 1999; Rodríguez-Caparrós et al, 2022) and is therefore consistent with regulation mediated by the enhancer. In agreement with this hypothesis, *Cdh24* transcript levels were elevated in DP thymocytes from *Rag2*^-/-^ x TCRβtg mice compared with DN3 thymocytes from *Rag2*^-/-^ mice (Fig. 6C). To directly assess the dependency of *Cdh24* transcription on Eα during thymocyte development, we analyzed DN3 and DP thymocytes derived from PBS- and anti-CD3ε-injected *Rag2*^-/-^, *Rag2*^-/-^ x Eα+EαCBE^-/-^, and *Rag2*^-/-^ x Eα^-/-^ mice (Fig. 6D). Consistent with a previous report (Rodríguez-Caparrós et al, 2022), *Cdh24* transcription was dependent on Eα+EαCBE in DP thymocytes, but was independent in DN3 thymocytes and non-T lineage tissues (Figs. 6D and S9B). Importantly, our data further demonstrate that *Cdh24* transcription in DP thymocytes depends on Eα even when the EαCBEs remain intact (Fig. 6D). These findings sharply contrast with the results obtained for *Dad1* and *Haus4* (Figs. 5 and S8), indicating that the EαCBEs are unable to insulate the distal *Cdh24* gene from enhancer activity. Thus, whereas Eα fails to overcome the CTCF-mediated barrier to regulate *Dad1* and *Haus4*, it successfully drives *Cdh24* transcription despite the presence of the EαCBEs.

**Figure 6.**
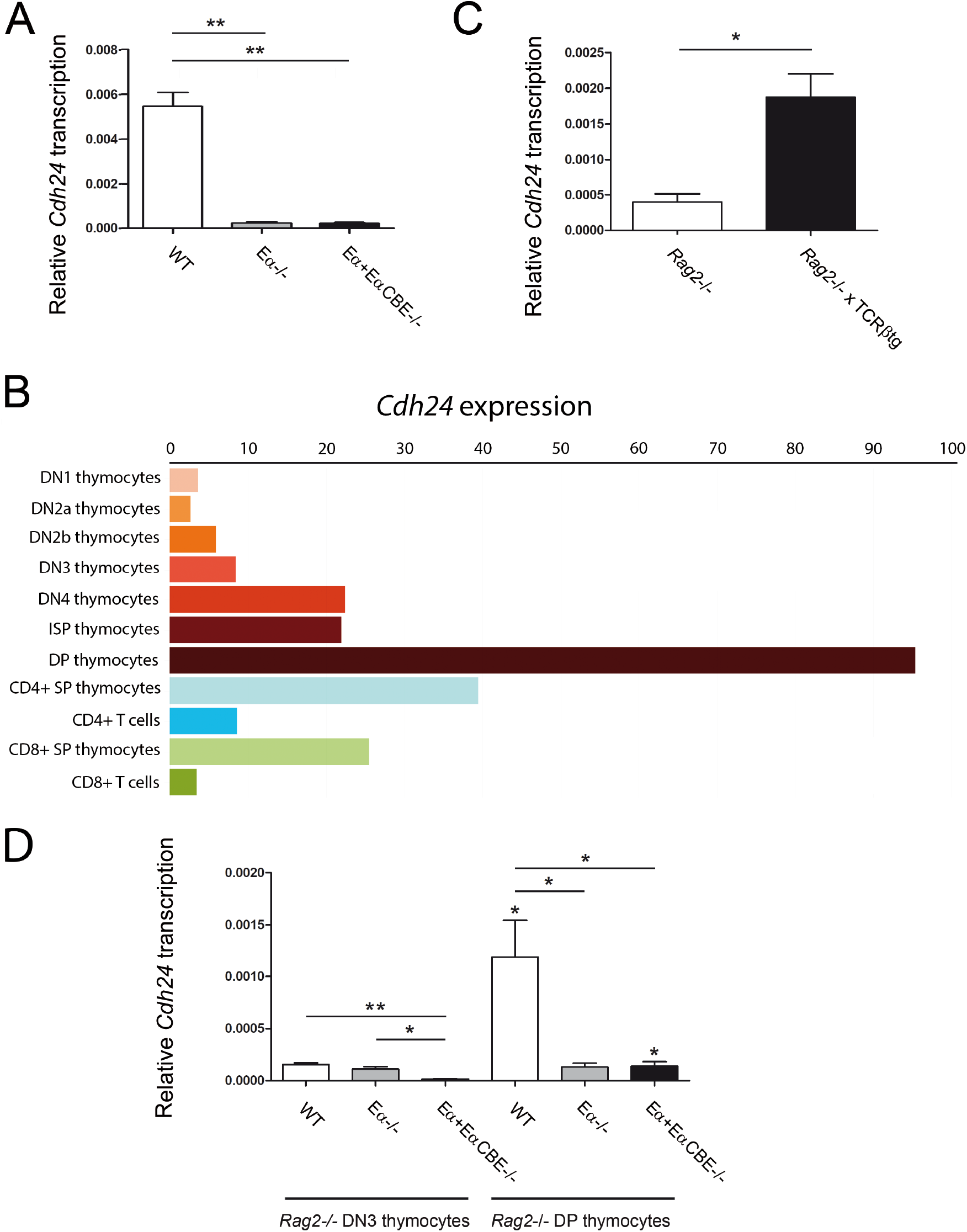
Eα-dependent transcription of *Cdh24*. **(A)** RT-qPCR analysis of *Cdh24* transcript levels in thymocytes from WT, Eα+EαCBE^-/-^, and Eα^-/-^ mice (n=2-4). Statistical analyses were performed using a non-parametric unpaired Student’s *t* test with Welch correction. Significance is indicated as follows: \**p* < 0.05, and \*\**p* < 0.005. **(B)** Expression profiles of *Cdh24* across thymocyte and mature T-cell subsets derived from the Immgen database; expression values were normalized using DESeq. **(C)** RT-qPCR analysis of *Cdh24* transcript levels in thymocytes from *Rag2*^-/-^ and *Rag2*^-/-^ x TCRβtg mice (n=3). **(D)** RT-qPCR analysis of *Cdh24* transcript levels in PBS-injected or anti-CD3ε-injected *Rag2*^-/-^, *Rag2*^-/-^ x Eα^-/-^, and *Rag2*^-/-^ x Eα+EαCBE^-/-^ mice (n=3-7). Transcript levels were normalized to *Actb* and presented as mean ± SEM from duplicate RT-qPCR measurements across the indicated number (n) of independent experiments. Statistical analyses were performed using a non-parametric unpaired Student’s *t* test with Welch correction. Significance is indicated as follows: \**p* < 0.05, and \*\**p* < 0.005.

### Genomic interactions between Eα+EαCBE and regions within the telomeric TAD

Previous Hi-C analyses revealed prominent long-range interactions between the Eα+EαCBE region and the telomeric TAD (Seitan et al, 2013; Zhao et al, 2020). Consistent with these observations, 4C-seq analyses using either the Eα+EαCBE HindIII fragment or the EαCBE DpnII fragment as viewpoints (Rodríguez-Caparrós et al, 2022) detected low-frequency interactions with the GRCm38/mm10_chr14:54642250-54685768 region (dark blue-to-orange fragments) or the GRCm38/mm10_chr14:54637759-54645134 region (turquoise-to-dark green fragments), respectively (Figs. 7A and 8). These interacting regions encompass the *Cdh24* promoter, which contains a CBE (CTCF peak 6) oriented opposite to the two EαCBEs (CTCF peak 3), as evidenced by interactions involving the dark blue and green fragments (Figs. 8A and 8B), and the turquoise-to-green fragments (Figs. 8C and 8D). In addition to these contacts, other regions within *Acin1* also exhibited robust interactions, predominantly with the Eα+EαCBE fragment, as indicated by interactions involving the pink and orange fragments (Figs. 8A and 8B), but also with the EαCBE fragment, as evidenced by interactions with pink-to-dark green fragments (Figs. 8C and 8D). The interacting *Acin1* region includes three CTCF peaks (peaks 7-9) comprising four CBEs, two of which, those associated with peaks 8 and 9, are oriented oppositely to the two EαCBEs (Figs. 7B-7D). Interactions within the telomeric TAD were further examined by 3C analyses performed in DN3 and DP thymocytes using the Eα+EαCBE HindIII fragment as the anchor (Fig. 9 and Table S1). Restriction enzyme digestion and ligation efficiencies were comparable between DN3 and DP samples (Fig. S10). As previously reported (del Blanco et al, 2015; Rodríguez-Caparrós et al, 2022; Shih et al, 2012), strong interactions between the Eα+EαCBE region and the T early α exon promoter (TEAp), located 68.6 kb upstream of the centromeric end of the anchor fragment, were selectively observed in DP thymocytes. By contrast, no interaction was detected with a nearby region located 72.3 kb upstream, previously referred to as the “-75-bp region”, which lies 591 bp downstream of *Trdv5* and 5,054 bp upstream of *Trav61* (Fig. 9). In agreement with the HindIII 4C-data shown in Fig. 7A, we observed a distance-dependent gradient of progressively weaker yet robust interactions between the Eα+EαCBE fragment and the downstream genes *Dad1, Haus4*, and *Cdh24*, located +12.9, +325.6, and +404.6, respectively, relative to the 3’ end of the anchor fragment. In contrast, no interactions were detected with distal control regions at *Dhrs2*, located +1010.3 and +1011.2 kb from the anchor (Fig. 9A). Importantly, all detected interactions were developmentally regulated and were either absent or markedly reduced in DN3 thymocytes compared with DP thymocytes (Fig. 9A). Notably, and consistent with the 4C data, a 17-kb region containing *Acin1* sequences located +450.7 kb downstream of the anchor exhibited a strong interaction with the Eα+EαCBE region (Figs. 8A, 8B, and 9). These results confirm that Eα+EαCBEs strongly interacts with an *Acin1* region positioned between CTCF peaks 8 and 9 that harbors two CBEs oriented oppositely relative to the EαCBEs (Figs. 7 and 9). Despite these physical interactions, Eα does not regulate *Acin1* transcription, as its regulatory activity within the telomeric TAD is restricted to *Cdh24* (Figs. 6 and S11). Although our 3C experiments failed to detect specific physical interactions between the Eα+EαCBE region and the *Cdh24* promoter, this apparent discrepancy relative to the 4C-seq results is most likely attributable to the greater sensitivity of 4C-seq compared with 3C for detecting weak or low-abundance genomic interactions. Collectively, these findings demonstrate that Eα-dependent regulation of *Cdh24* involves functional inter-TAD physical contacts between the Eα+EαCBE region and the telomeric TAD, enabling productive E-P interactions and Eα-dependent activation of the *Cdh24* promoter. These findings underscore the ability of a strong enhancer positioned at a TAD boundary to bypass topological insulation in a specific T-cell developmental context.

**Figure 7.**
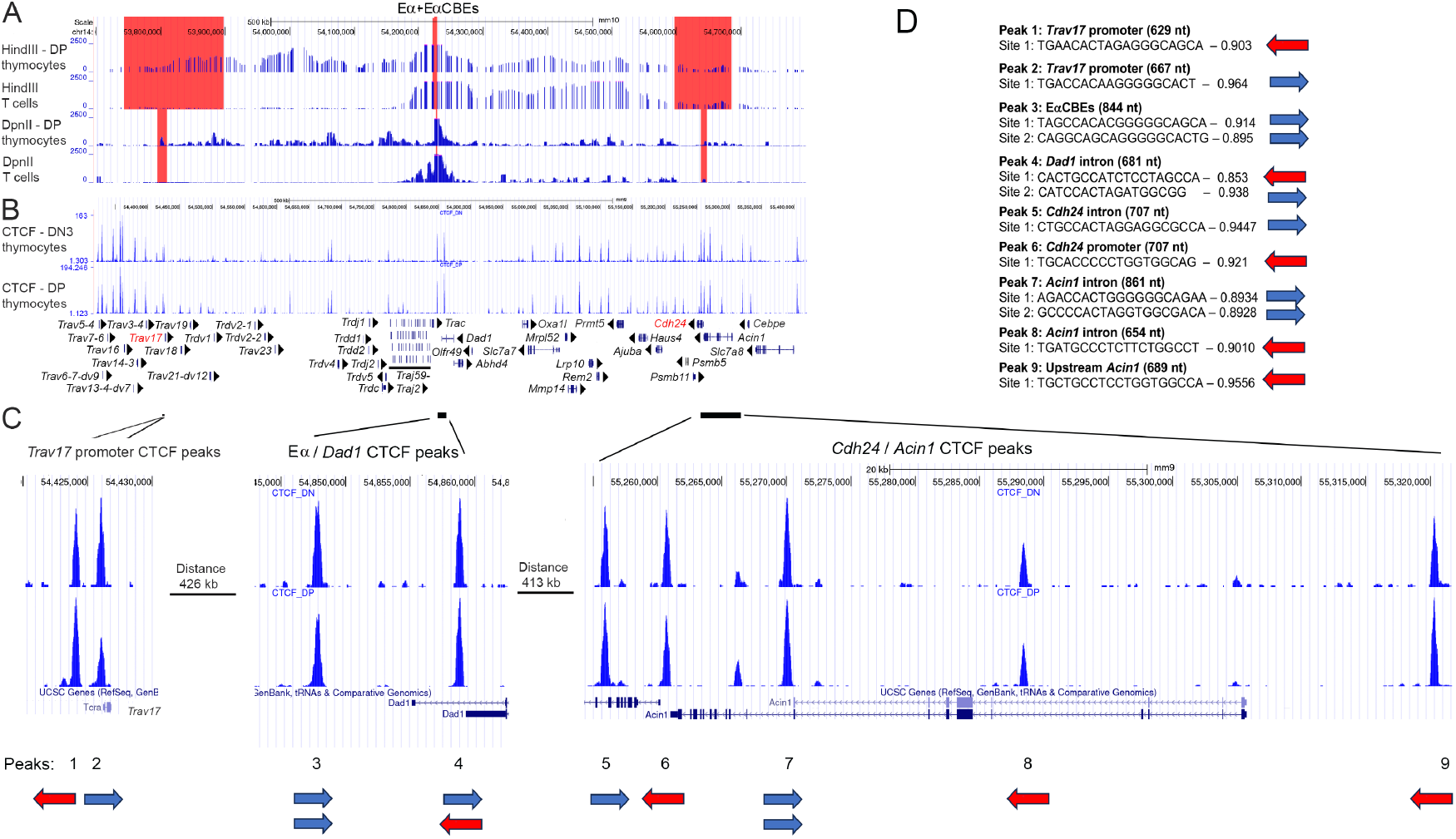
Long-range genomic contacts involving Eα+EαCBEs or EαCBE and CTCF binding within the centromeric and telomeric TADs. **(A)** 4C-seq interaction profiles generated using the Eα+CBE HindIII fragment or EαCBEs DpnII fragment viewpoints (highlighted in red) at the unrearranged locus in DP thymocytes from anti-CD3ε-treated *Rag2*^-/-^ mice and at the rearranged locus in WT T cells. Results are representative of two independent experiments. Data are shown as reads per million mapped reads and aligned to the GRCm38/mm10 mouse reference genome using the UCSC Genome Browser. **(B)** CTCF binding assessed by ChIP-seq at the unrearranged locus in DN3 and DP *Rag2*^-/-^ thymocytes. Data are shown as reads per million mapped reads and aligned to the NCBI37/mm9 mouse reference genome using the UCSC Genome Browser. **(C)** Close-up view of CTCF binding at the *Trav17* promoter, EαCBEs, and the *Cdh24*/*Acin1* region. **(D)** Schematic representation of CBEs and their orientations. Genomic coordinates (GRCm39/mm39) of the CBEs are listed in Table S3.

**Figure 8.**
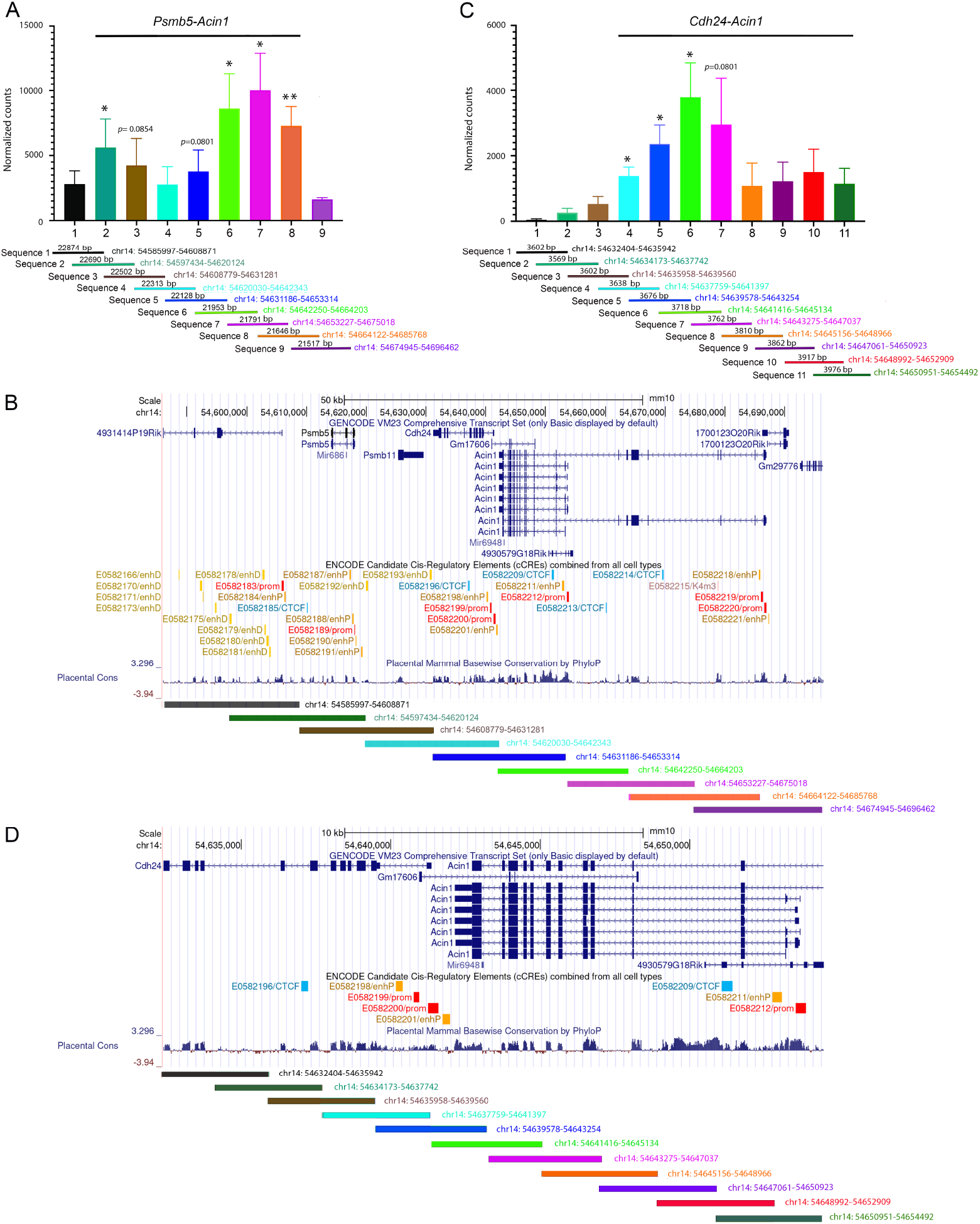
Interactions between EαCBEs and genomic regions. **(A)** 4C-seq experiments were performed in DP thymocytes from anti-CD3ε-treated *Rag2*^-/-^ mice and in WT T cells using the Eα+EαCBE HindIII fragment as the viewpoint. Interaction signals were normalized using 4C-ker and combined. Significant interactions between Eα+EαCBEs and multiple overlapping genomic fragments were detected. Data are presented as mean ± SEM from two independent experiments, combining both sample types. Statistical significance was assessed using a non-parametric unpaired Student’s test with Welch correction, with fragment 1 used as reference: \**p* < 0.05, and \*\**p* < 0.005. **(B)** Genomic region spanning *Psmb5* to *Acin1* highlighting the significant interactions detected in A. **(C)** 4C-seq experiments were performed in DP thymocytes from anti-CD3ε-treated *Rag2*^-/-^ mice and in WT T cells using the EαCBE DpnII fragment as viewpoint. Signals were normalized using 4C-ker and combined. Significant interactions were detected between the EαCBEs and multiple overlapping genomic fragments, across a region extending from the telomeric end of *Cdh24* to the centromeric end of *Acin1*, as indicated by overlapping turquoise, blue, and bright green fragments. Data are presented as mean ± SEM from two independent experiments, combining both sample types. Statistical significance was assessed using a non-parametric unpaired Student’s test with Welch correction, with fragment 1 used as reference: \**p* < 0.05. **(D)** Genomic view of the *Cdh24/Acin1* region highlighting the significant interactions detected in panel C. Genomic regions are mapped to the mm10 mouse reference genome using the UCSC Genome Browser. Candidate cis-Regulatory Elements derived from the ENCODE and placental conservation data are indicated.

**Figure 9.**
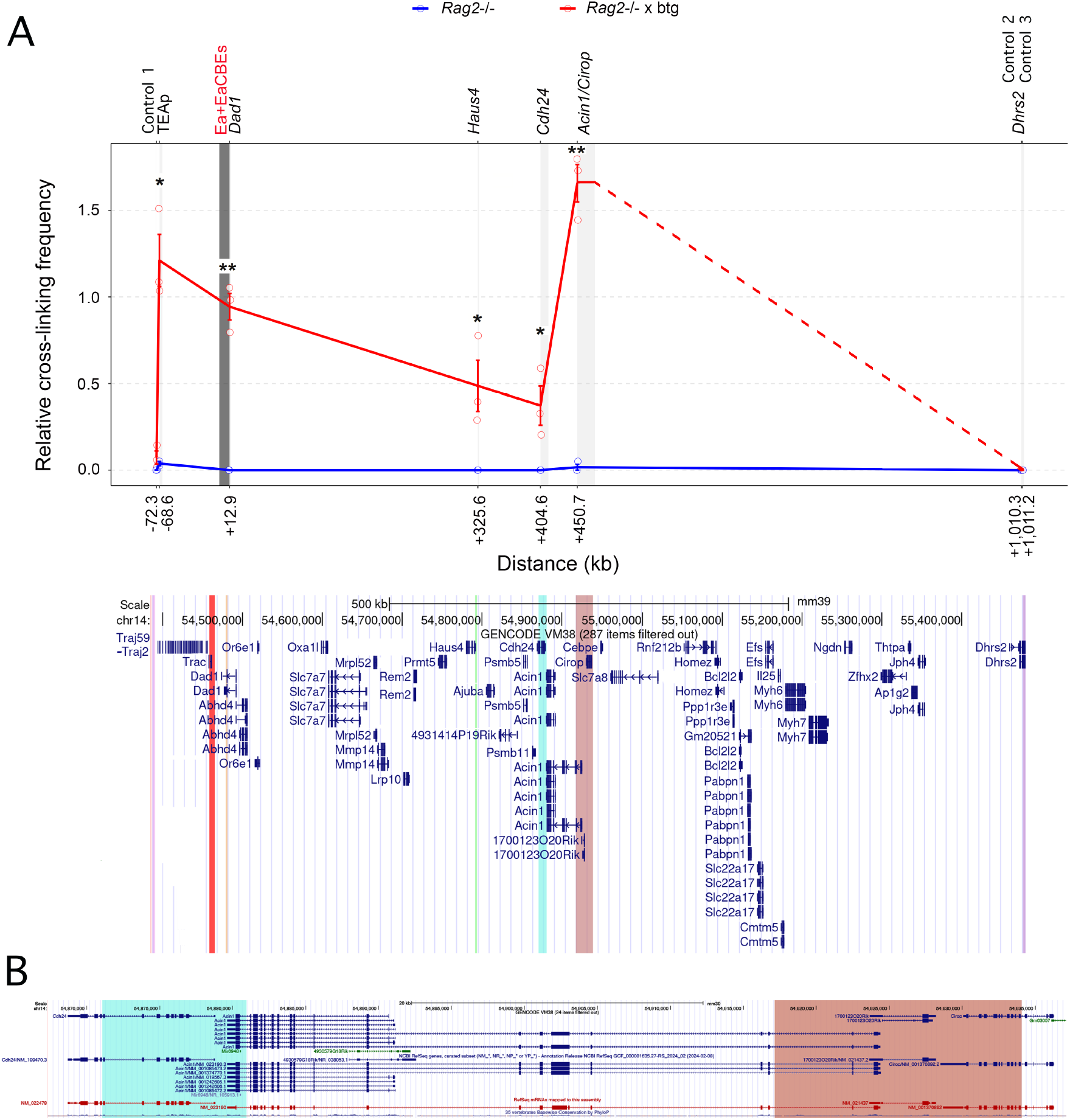
Developmentally regulated interactions between Eα+EαCBEs and regions within the centromeric and telomeric TADs. **(A)** Chromatin contacts between Eα+EαCBEs (region highlighted in red) and multiple were assessed by 3C analysis in DN3 (blue line) and DP (red line) thymocytes. Tested regions include a control fragment located 72.3 kb upstream of Eα (Control 1), the TEA α promoter (TEAp; highlighted in pink), *Dad1* (salmon), *Haus4* (green), *Cdh24* (turquoise), *Acin1/Cirop* (brown), and two distal control regions located more than 1 Mb away at *Dhrs2* (Controls 2 and 3, purple). Genomic regions are mapped to the GRCm39/mm39 mouse reference genome using the UCSC Genome Browser. Genomic coordinates of the analyzed fragments are listed in Table S1). Interaction frequencies were normalized to the ligation product between Eα+EαCBE-containing HindIII fragment and a HindIII fragment located 8.9 kb upstream of the enhancer. Data are shown as mean ± SEM of duplicate qPCRs from two independent DN3 samples and three independent DP samples. Statistical comparisons between DN3 and DP thymocytes were performed using a non-parametric unpaired Student’s *t* test with Welch correction. Statistical significance was assessed using a non-parametric unpaired Student’s test with Welch correction, with control fragments used as reference: \**p* < 0.05, and \*\**p* < 0.005. **(B)** Enlarged view of the genomic region spaning *Cdh24* to *Cirop*, highlighting the analyzed *Cdh24* fragment (turquoise) and the *Acin/Cirop* fragment (brown).

## Discussion

Hi-C analyses have revealed that genomic regions within chromosome territories exhibit strong self-association, forming higher-order structures known as TADs. These domains are typically limited by convergently oriented CBEs and are established through cohesin-dependent loop extrusion, during which chromatin is threaded through cohesion complexes anchored by CTCF at CBEs. Within TADs, dynamic chromatin looping facilitates long-range E-P interactions, enabling precise regulation of gene expression across large genomic distances (da Costa Nunes & Noordermeer, 2023). However, accumulating evidence challenges the view that TAD boundaries strictly insulate functional E-P interactions or are universally required for transcriptional control (20). In multiple instances, gene expression remains largely unaffected following chromosomal rearrangements that disrupt TAD boundaries (Despang et al, 2019; Dong et al, 2018; Ghavi-Helm et al, 2019; Laugsch et al, 2019), and depletion of boundary-forming complexes, resulting in the loss of TAD architecture, often leads to only modest transcriptional effects (Hsieh et al, 2022; Nora et al, 2017; Rao et al, 2017; Schwarzer et al, 2017; Soshsnikova et al, 2017). Notably, approximately one-third of long-range E-P interactions identified in mice, humans, and *Drosophila* span TAD boundaries (Balasubramanian et al, 2024; Freire-Pritchett et al, 2017; Galupa & Crocker, 2020; Galupa et al, 2020; Hsieh et al, 2022; Javierre et al, 2016). Despite their prevalence, the functional significance of most of these inter-TAD contacts remains uncertain, as a substantial fraction occurs in contexts where the target gene is not actively transcribed (Espinola et al, 2021; Ghavi-Helm et al, 2014; Ing-Simmons et al, 2021). Whether such interactions represent transcriptionally inert background contacts or contribute to higher-order genome organization remains an open question. Nevertheless, functional inter-TAD E-P interactions have been clearly demonstrated in specific biological contexts. These include the *twist* locus in *Drosophila*, revealed through enhancer relocation experiments (Balasubramanian et al, 2024), as well as several mammalian examples involving developmentally regulated enhancers controlling limb and craniofacial patterning genes that are separated from their target genes by TAD boundaries (Beccari et al, 2021; Hung et al, 2024; Kessler et al, 2023; Rouco et al, 2021). In this study, we show that Eα-dependent activation of *Cdh24* transcription in mouse thymocytes constitutes a compelling example of a naturally occurring long-range inter-TAD E-P interaction in an adult mammalian context during T-cell development. Together, these findings support the notion that, under specific developmental conditions, enhancers positioned at TAD boundaries can overcome topological constraints to regulate distal gene expression.

Chromosome conformation capture-based approaches have revealed extensive long-distance interactions between Eα+EαCBEs and genomic regions within both the adjacent centromeric and telomeric TADs in *Rag2*^-/-^ DN3 and DP thymocytes, as well as in WT T cells (Carico & Krangel, 2015; Chen et al, 2015; Dai et al, 2023; Rodríguez-Caparrós et al, 2022; Seitan et al, 2013; Shih et al, 2012; Zhao et al, 2020), as illustrated by the 4C-seq data in Figs. 7, 8, and S12. Interactions extending from the EαCBEs to *Trav17* define the centromeric TAD, whereas interactions reaching *Cdh24* delineate the telomeric TAD (Carico et al, 2017; Dai et al, 2023; Rodríguez-Caparrós et al, 2022; Shih et al, 2012; Zhao et al, 2020), as shown in Figs. 1, 7, and S12. The upstream boundary of the centromeric TAD is defined by two CTCF-binding peaks located at the *Trav17* promoter: an upstream peak (CTCF peak 1) containing a CBE oriented upstream, and a downstream peak (CTCF peak 2) containing a CBE oriented downstream (Fig. 7B-7D). On the other hand, the downstream boundary of the telomeric TAD is defined by five CTCF-binding peaks (CTCF peaks 5-9) located within the *Cdh24* promoter/*Acin1* region, comprising six CBEs, three of which are oriented opposite to the EαCBEs (Figs. 7B-7D). Although these domains have previously been referred to as sub-TADs (Rodríguez-Caparrós et al, 2022; Seitan et al, 2013; Zhao et al, 2020), this distinction appears largely semantic. Given their genomic size, tissue-specific activity, conservation across cell types, and essential roles in cell differentiation and gene regulation (Zhao et al, 2020), these domains fulfill the criteria of bona fide TADs.

Previous analyses of mice lacking both Eα and the associated EαCBEs revealed a marked reduction in gene transcription across both TADs in DP thymocytes, as well as significantly decreased transcription of *Dad1* and *Haus4* in DN3 thymocytes and non-T lineage tissues (Rodríguez-Caparrós et al, 2022; Sleckman et al, 1997; Zhao et al, 2020). This reduction in telomeric TAD gene transcription was interpreted as a consequence of structural disruption resulting from the loss of the EαCBEs (Rodríguez-Caparrós et al, 2022; Zhao et al, 2020). Consistent with a role for EαCBEs in delimiting the boundary between the flanking TADs, 4C-seq experiments demonstrated that the centromeric EαCBE preferentially interacted with sequences within the centromeric TAD, whereas the telomeric EαCBE preferentially interacted with sequences in the telomeric TAD (Zhao et al, 2020). Moreover, specific deletion of the two EαCBEs while preserving Eα affected transcription in both domains in DP thymocytes (Zhao et al, 2020). In the centromeric TAD, EαCBE deletion impaired primary *Tcra* rearrangements by reducing accessibility of proximal *Trav* gene segments, likely reflecting disrupted contacts between EαCBEs and CBEs associated with proximal *Trav* promoters. In the telomeric TAD, deletion of EαCBE resulted in a modest increase in transcription of the proximal genes *Dad1* and *Abdh4*, accompanied by a slight decrease in transcription of the more distal genes *Ajuba* and *Cdh24*. These observations suggest that EαCBEs limit the positive influence of Eα on proximal genes within the telomeric TAD, while potentially facilitating enhancer interactions with more distal targets such as *Cdh24*.

To directly assess the role of Eα in transcriptional regulation across the flanking TADs, we generated Eα^-/-^ mice in which the EαCBEs were preserved. Deletion of Eα alone is not expected to perturb global three-dimensional genome organization, since removal of the two EαCBEs does not disrupt TAD formation (Zhao et al, 2020). Our data demonstrate that the deleted Eα region contains the essential regulatory sequences required for *Tcra-Tcrd* transcription and TCRαβ expression in thymocytes and mature αβ T lymphocytes. Importantly, in the presence of intact EαCBEs, Eα failed to activate transcription of the proximal telomeric TAD genes *Dad1* and *Haus4*, but efficiently activated transcription of the distal telomeric TAD gene *Cdh24*. Independent support to these findings comes from Eα E-box mutant mice (Mihai et al, 2023), which exhibit normal transcription of *Dad1, Abdh4, Prmt5*, and *Ajuba* but reduced *Cdh24* transcription. Thus, two independently generated CRISPR/Cas9 mutant models, Eα^-/-^ and EαE-box^-/-^, displayed identical phenotype: preserved transcription of proximal telomeric TAD accompanied by reduced *Cdh24* transcription. Together, these results indicate that Eα drives transcription within the centromeric TAD (*Tcra-Tcrd*) and selectively activates *Cdh24* at the distal end of the telomeric TAD, while remaining constrained from regulating proximal telomeric genes by the intervening EαCBEs. This supports a model in which EαCBEs function as an insulating boundary that separates Eα from proximal telomeric promoters while permitting E-P interactions with the distal genes (Zhao et al, 2020). Our data further show that the strong interactions detected between the Eα+CBE region and the *Dad1* and *Haus4* genes by 4C-seq and 3C analysis ((Rodríguez-Caparrós et al, 2022), and Figs. 7 and 9), as well as by Hi-C (Seitan et al, 2013), do not result in transcriptional activation. These observations are consistent with previous reports describing abundant non-functional E-P interactions that span TAD boundaries without measurable effects on gene expression (Freire-Pritchett et al, 2017; Galupa & Crocker, 2020; Galupa et al, 2020; Hsieh et al, 2022; Javierre et al, 2016). Because Eα does not activate *Dad1* and *Haus4* transcription, additional regulatory elements may contribute to repression of these genes in Eα+EαCBE^-/-^ tissues, potentially acting as silencers whose activity becomes apparent in the absence of Eα (Hussain et al, 2023). Notably, *Cdh24* is expressed at very low levels in most immune cell type but is highly expressed in DP thymocytes and specific subsets of γδ T-cells (www.immgen.org). By analogy with the role of low-affinity interactions mediated by MHC-presented peptides and E-cadherin during immunological synapse formation and spindle organization at β-selection (Charnley et al, 2023; Irving et al, 1998; Mallis et al, 2015), Eα-dependent *Cdh24* expression in DP thymocytes may enhance interactions between the stromal cells and DP thymocytes, thereby promoting synapse formation during positive selection. Thus, Eα-mediated regulation of *Cdh24* expression may complement its essential role in driving TCRαβ expression during T cell development.

Our analysis of a targeted enhancer loss-of-function mutant in defined T cell populations provides a powerful framework to dissect mechanisms by which long-range inter-TAD E-P interactions influence chromatin structure and transcriptional activation. Our findings align with the emerging concept that enhancers positioned at TAD boundaries possess a unique ability to establish functional inter-TAD interactions with distal regions located at the edges of neighbouring domains, thereby coordinating transcriptional regulation across adjacent TADs (Smith et al, 2016). Recent molecular dynamics simulations of *Pitx1* regulation by the Pen enhancer have proposed a “stacked-border” model, in which TAD boundaries cluster into a shared hub to facilitate interactions between boundary-associated enhancers and promoters (Hung et al, 2024). These stacking interactions likely arise through loop-extrusion and may contribute to the formation of architectural stripes observed in Hi-C maps. Although the molecular mechanisms underlying functional inter-TAD enhancer-dependent activation of promoters remain unclear, stacked borders formed between the EαCBEs and the *Acin1*-interacting region may facilitate functional communication between Eα and the *Cdh24* promoter. The physical interactions between the EαCBEs and *Acin* sequences may involve the two inversely oriented CBEs (CTCF peaks 8 and 9) that flank the *Acin1* HindIII fragment, which exhibits strong interactions with the Eα+EαCBE region in our 3C and 4C-seq analyses (Figs. 7-9). This spatial arrangement may promote the formation of transcriptional condensates enriched in RNA polymerase II, Mediator, TFs, and additional cofactors, thereby enabling transcriptional activation of specific genes (Cho et al, 2016; Cho et al, 2018; Cisse et al, 2013; Du et al, 2024; Iborra et al, 1996; Li et al, 2019; Wang et al, 2022). Such condensates can dynamically position enhancers and promoters within submicron proximity, enabling transcriptional bursting (Du et al, 2024). The ability of enhancers and promoters separated by hundreds of nanometers to communicate through these condensates, together with chromatin motion at the ∼200 nm scale that is sufficiently rapid to permit frequent transient contacts, blurs the distinction between direct physical contact and action-at-distance models of gene regulation (Mazzocca et al, 2025; Yang & Hansen, 2024). Within this framework, the proximity of both Eα and the *Cdh24* promoter to a TAD boundary capable of mediating chromatin loops may facilitate transient E-P interactions within transcriptionally active condensates, thereby driving developmentally regulated gene expression.

## Methods

### Mice

The *Rag2*^-/-^, *Rag2*^-/-^ x TCRβtg, Eα+EαCBE ^-/-^, *Rag2*^-/-^ x Eα+EαCBE^-/-^, and *Rag2*^-/-^ x TCRβtg x Eα+EαCBE^-/-^ mice have been described previously (Hernández-Munain et al, 1999; Shinkai et al, 1992; Sleckman et al, 1997). The Eα^-/-^ mice were generated by CRISPR/Cas9-mediated genome editing in C57bl/6 embryonic stem cells at the Transgenesis and Genetic Edition Service at CNB-CSIC, Madrid, Spain. The 200-bp Eα region, containing the Tα1-Tα4 elements, was deleted using Alt-R-CRISPR-Cas9 crRNAs (IDT) containing target-specific protospacer sequences (listed in Table S2), along with a 16-nucleotide tracrRNA fusion domain. These guide RNAs included proprietary chemical modifications to protect against RNase degradation. The crRNAs were duplexed with the 67-nucleotide Alt-R tracrRNA (IDT) to form functional crRNA:tracrRNA duplexes and used in combination with Alt-R HiFi Cas9 nuclease (IDT). Two independent founder lines were selected: 13094 (Eα1^+/-,^ referred to as the 700 strain) and 13097 (Eα2^+/-^, referred to as the 800 strain). Founders were outcrossed to WT C57bl/6 mice (Charles River, Spain) for three generations prior to establishing homozygous Eα1^-/-^ and Eα2^-/-^ lines. The deleted sequences and guide RNA positions of both lines are shown (Figs. S2 and S3). *Rag2*^-/-^ x Eα1^-/-^ mice (referred to as the 900 mice) were generated by crossing Eα1^-/-^ animals with C57bl/6 *Rag2*^-/-^ mice. All experiments were conducted using adult mice (4-10 weeks old). Animals were maintained under pathogen-free conditions in the Animal Experimentation Unit of the IPBLN-CSIC.

### Cell and tissue preparations

Thymocytes and splenocytes were obtained by mechanical dissociation of the thymus and spleen through 70-μm nylon mesh strainers (Pluriselect, 43-50070-51). Single-cell suspensions from liver, kidney, and testis extracts were prepared using a gentleMACS Dissociator (Miltenyi Biotech) according to the manufacturer’s instructions.

### *In vivo* induction of DN3 to DP differentiation

To induce DN3 to DP thymocyte differentiation, adult *Rag2*^-/-^, *Rag2*^-/-^ x Eα+EαCBE ^-/-^, and *Rag2*^-/-^ x Eα^-/-^ mice, animals were intraperitoneally injected with 150 μg of anti-CD3ε antibody (clone 145-2C11) and euthanized 10 days later, as previously described (Rodríguez-Caparrós et al, 2019). Control mice received intraperitoneal injections of phosphate-buffered saline (PBS).

### Flow Cytometry-based cell analyses and separation

Total thymocytes and splenocytes from WT, Eα+EαCBE^-/-^, and Eα^-/-^ mice were stained with the following fluorochrome-conjugated antibodies: anti-CD4-FITC (clone GK1.5), anti-CD8α-PE (clone 53-6.7), anti-CD3ε-BV421 (clone 145-2C11), anti-TCRβ-FITC (clone H57-597), anti-TCRγδ-PE (clone GL3) and/or anti-Vα2-PE (clone B20.1). All antibodies were obtained from Biolegend. Samples were analyzed using a FACSymphony flow cytometer (BD Biosciences). For γδ T cell purification, splenocytes were stained with anti-CD3ε-BV421 (clone 145-2C11) and anti-TCRγδ-PE (clone GL3), and double-positive stained cells were isolated using a FACSAria-III cell sorter (BD Biosciences).

### Western blot

Cells and tissue extracts were lysed in RIPA buffer containing 50 mM Tris-HCl (pH 7.5), 150 mM NaCl, 1% NP-40, 0.5% sodium deoxycholate, 0.05% SDS, 1 mM EDTA, 1 mM dithiothreitol, 1 mM PMSF, and a protease inhibitor cocktail (Complete, Roche). Lysates were incubated for 30 min at 4°C and centrifuged at 16,000 x g for 5 minutes at 4°C. Protein quantification was quantified using the detergent-compatible colorimetric DC Protein Assay Kit (Bio-Rad), and 500μg of total protein was resolved by SDS-PAGE using 1.5 mm, 15% SDS polyacrylamide gels. Proteins were transferred onto nitrocellulose membranes (Amersham Biosciences) and incubated with the following primary antibodies at 4°C for 20-24 hours: anti-DAD1 (EpigenTEK, A70728; 2 μg/mL), anti-HAUS4 (FineTest, FNab01027; 1 μg/mL), and anti-GAPDH (Santa Cruz Biotechnology, sc-365062; 0.4 μg/mL). After washing, membranes were incubated with horseradish peroxidase-conjugated secondary antibodies: goat anti-rabbit IgG (Abcam, AB205718; 1:5000) for anti-DAD1 and anti-HAUS4, or goat anti-mouse IgG-HRP (Agilent Dako, P044701; 1:500) for anti-GAPDH. Signal detection was performed using enhanced chemiluminiscence reagents (Cytiva-Amersham), and membranes were imaged using a Curix 60 film developer (Agfa).

### Quantitative RT-PCR (RT-qPCR)

Total RNA was extracted using TRIdity reagent (Panreac AppliChem) following the manufacturer’s instructions. First-strand cDNA was synthesized from 500 ng of total RNA using PrimeScript RT Master Mix (RR036, Takara) in a final volume of 20 µL, and subsequently diluted to 100 µL with Milli-Q water. Quantitative PCR (qPCR) was performed using 4 µL of diluted cDNA in a total volume of 10 µL per reaction, in duplicate, with the TB Green Premix Ex TaqII kit (RR820, Takara) on a Bio-Rad CFX-96 System. PCR cycling conditions were as follows: initial denaturation at 95°C for 7 minutes; 40 cycles of 95°C for 30 seconds, 59.5°C for 45 seconds, and 72°C for 30 seconds; followed by a final incubation at 95°C for 1 minute. Melting curve analyses were performed from 55°C to 90°C in 0.5°C increments (5 seconds per step) to verify specificity and the presence of as single amplicon. Relative gene expression was calculated using the ΔCt method and normalized to *Actb* transcription. Primers used were previously described (del Blanco et al, 2015; Rodríguez-Caparrós et al, 2019) and synthesized by Metabion, and are listed in Table S2.

### Quantitative chromatin immunoprecipitation (ChIP-qPCR)

ChIP-qPCR experiments were performed using chromatin from 2×10^7^ thymocytes. Chromatin was incubated with 5 µg of anti-CTCF antibody (Millipore, 07-729) or control (Abcam, ab46540) antibody, following previously described protocol (del Blanco et al, 2015). qPCR conditions used are described above. Primers specific for the EαCBEs region were previously reported (del Blanco et al, 2015) and synthesized by Metabion. Primer pairs mapping to an EαCBE-containing region have been previously described (del Blanco et al, 2015; Rodríguez-Caparrós et al, 2019). In addition, primers mapping a region downstream of the EαCBEs present in WT, Eα ^-/-^, and Eα+EαCBE^-/-^ mice were synthesized by Metabion and are listed in Table S2.

### Chromosome conformation capture (3C) experiments

3C experiments were performed following a previously described protocol (Krijger et al, 2019; Rodríguez-Caparrós et al, 2022). Briefly, 10^7^ DN3 thymocytes from *Rag2*^-/-^ mice or DP thymocytes from *Rag2*^-/-^ x TCRβtg mice were resuspended in 5 mL RPMI 1640 with 10% fetal calf serum (FCS) and crosslinked by adding 5 mL of freshly prepared 4% paraformaldehyde (Thermo Fisher Scientific, 28908) in RPMI 1640 containing 10% FCS (final concentration 2%). Samples were mixed and incubated for 10 minutes at room temperature on a tube roller. Crosslinking was quenched with 1.5 mL of cold 1M glycine (final concentration 0.125 M), and cells were placed on ice. Thymocytes were centrifuged (7 minutes, 500 x g, 4°C), resuspended in 1 mL of cold Hank’s Balanced Salt Solution (HBSS), transferred to 1.5-mL tubes, centrifuged again (5 minutes, 500 x g, 4°C), and lysed in 1 mL of cold lysis buffer (150 mM NaCl, 50 mM Tris pH 7.5, 0.5% Nonidet P40 substitute (Fluka, 74385), 1% Triton X-100 (VWR, 28817.295), 5mM EDTA, and protease inhibitors (cOmplete Mini, Roche, 11836153001) for 20 minutes on ice. Nuclei were pelleted by centrifugation (5 minutes, 500 x g, 4°C), washed with 500 µL of 1.2X restriction enzyme buffer (New England Biolabs buffer #2), resuspended in 500 µL of 1.2X restriction enzyme buffer, and warmed to 37°C. SDS (0.3% final) were added and incubated for (1 hour, 750 rpm, 37°C), followed by Triton X-100 (2.5% final) for another 1-hour incubation. Chromatin was digested with 200 U of HindIII (New England Biolabs, RO104) (3 hours, 750 rpm, at 37°C), followed by additional 200 U of HindIII (overnight, 750 rpm, at 37°C). Digestion was assessed by agarose gel electrophoresis of control aliquots treated with proteinase K. After confirming digestion, HindIII was inactivated (20 minutes, 65°C), and samples were ligated in T4 DNA ligase buffer (66 mM Tris pH:7.5, 5 mM MgCl2, 5 mM DTT, and 1 mM ATP) in a volume of 7 mL (overnight, 16°C) using 4000 U of T4 DNA Ligase (New England Biolabs, M0202). Ligation efficiency was verified by gel electrophoresis of control aliquots. Samples were treated with proteinase K (86 ng/mL final concentration, overnight, 65°C). DNA was extracted using phenol, phenol/chloroform, and chloroform and then precipitated and precipitated (overnight, -20°C) using sodium acetate, isopropanol, and Glycoblue (Thermo Fisher Scientific, AM2515). DNA pellets were washed with cold 70% ethanol and resuspended in 200 µL milli-Q water. Digestion efficiency (≥ 60-70%) at specific restriction sites was confirmed assessed by Syb^R^-Green qPCR using TB Green Premix Ex TaqII kit (Takara, RR820) on a Bio-Rad CFX-96 System. Efficiency was calculated using the formula: % restriction = 100-200/2^((Ct_R_-Ct_C_)_digested_-(Ct_R_-Ct_C_)_undigested_ (Hagège et al, 2007). The names of the sequences analyzed, their genomic coordinates based on the mouse GRCm39/mm39 genome assembly, the size of HindIII-generated fragments, and their linear distances to the anchor site are listed in Table S1. An Eα fragment unaffected by HindIII was used as control template. Primer efficiencies were validated on undigested templates. Interaction frequencies with Eα+CBEs were normalized to Eα-control fragment crosslinking (normalization fragment) (Hagège et al, 2007). qPCRs were performed in duplicate using 4 µL of diluted DNA in 10µL reactions, using the qPCR program described above. Primers used were synthesized by Metabion and listed in Table S2.

### Statistical analysis

All statistical analyses were performed using GraphPad Prism version 5.0 (GraphPad software). In all cases, a minimum of three independent experiments, each performed with different biological samples, were conducted, and the number of experiments is indicated in the figure legends. Data are presented as mean ± standard error of mean (SEM). Statistical comparisons were made using a non-parametric unpaired Student’s t-test with Welch correction, as appropriate. Statistical significance is indicated by asterisks as follows: *p* <0.05(*), *p* < 0.005 (**), and *p*<0.0005 (***). The absence of an asterisk denotes a change that was not statistically significant relative to the control.

## Abbreviations

3C: chromosome conformation capture
Cα: *Trac*
CBE: CTCF binding element
Cδ: *Trdc*
ChIP: chromatin immunoprecipitation
E-P: enhancer-promoter
Eα: *Tcra* enhancer
EαCBEs: Eα-associated CBEs
Eα+EαCBE: Eα and associated EαCBEs
DN: CD4^-^CD8^-^ double negative
DP: CD4^+^CD8^+^ double positive
PBS: phosphate-buffered saline
MFI: mean fluorescence intensity
RT-qPCR: quantitative RT-PCR
SEM: standard error of mean
SP: CD4^+^ or CD8^+^ single positive
TAD: topologically associating domain
TCR: T-cell receptor
TF: transcription factor
WT: wild-type

## Data availability

4C data was submitted to the BioProject, National Center for Biotechnology Information under accession number PRJNA749915 (Rodríguez-Caparrós et al, 2022). CTCF ChIP-seq were downloaded from GEO database GSE41743 (Shih et al, 2012).

## Author contributions

A.R.-C., L.L.-C., and J.L.-R. performed the experiments; J.L.-R. and V.C.-R. conducted mouse genotyping; M.D.-R., S.J.-L., and A.C.-M. assisted with Western-blot analyses; E.A.-L. carried out 4C-ker analyses; L.C.T.-C. assisted with Fig. 9 preparation. C.S. contributed to project discussions; C.H.-M. was responsible for planning and supervising the research, writing and editing the manuscript, and preparing the figures.

## Acknowledgements

We thank Belén Pintado and Verónica Domínguez from the Transgenesis and Editing Edition Service at CNB-CSIC for generating of Eα^-/-^ mice; Clara Sánchez González and Beatriz Ruiz Entrena for their support in maintaining the mouse colony at the IPBLN-CSIC Animal Experimentation Unit; and María M. Pérez Sánchez-Cañete and Marco Apolo Pulpillo for assistance with flow-cytometry analysis.

This work was supported by grants from the Ministerio de Ciencia, Innovación y Universidades [PID2021-128720NB-I00 to C.H.-M. and PID2020-118859GB-I00 to C.S.], Junta de Andalucía [P20-01271 to C.H.-M. and P20-01269 to C.S.], and Consejo Superior de Investigaciones Científicas [2025AEP113 to C.H.-M. and 2024AEP094 to C.S.]. Additional support was provided by the European Region Development Fund.

## References

Aifantis I, Bassing CH, Garbe AI, Sawai K, Alt FW, von Boehmer H (2006) The Eδ enhancer controls the generation of CD4-CD8-α?TCR-expressing T cells that can give rise to different lineages of α? T cells. J Ex Med 203: 1543–1550

Balasubramanian D, Borges-Pinto P, Grasso A, Vicent S, Tarayre H, Lajoignie D, Ghavi-Helm Y (2024) Enhancer-promoter interactions can form independently of genomic distance and be functional across boundaries. Nucleic Acids Res 52: 1702–1719

Beccari L, Jaquier G, Lopez-Delisle L, Rodríguez-Carballo E, Mascrez B, Gitto S, Woltering J, Duboule D (2021) Dbx2 regulation in limbs suggest interTAD sharing of enhancers. Dev Dyn 250: 1280–1299

Boudil A, Matei IR, Shih HY, Bogdanoski G, Yuan JS, Chang SG, Montpellier B, Kowalski PE, Voisin V, Bashir S, Bader GD, Krangel MS, Guidos CJ (2015) IL-7 coordinates proliferation, differentiation and Tcra recombination during thymocyte ?-selection. Nat Immunol 16: 397–405

Brewster JL, Martin SL, Toms J, Goss D, Wang K, Zachrone K, Davis A, Carlson G, Hood L, Coffin JD (2000) Deletion of Dad1 in mice induces an apoptosis-associated embryonic death. Genesis 26: 271–278

Busslinger GA, Stocsits RR, van der Lelig P, Axelsson E, Tedeschi A, Galjart N, Peters JM (2017) Cohesin is positioned in mammalian genomes by transcription, CTCF and Wapl. Nature 544: 503–507

Carico Z, Choudhury KR, Zhang B, Zhuang Y, Krangel MS (2017) Tcrd rearrangements redirects a processive Tcra recombination program to expand the Tcra repertoire. Cell Rep 19: 2157–2173

Carico Z, Krangel MS (2015) Chromatin dynamics and development of the TCRα and TCR8 repertories. Adv Immunol 128: 307–361

Charnley M, Allam AH, Newton LM, Humbert PO, Russell SM (2023) E-cadherin in developing murine T cells controls spindle alignment and progression during ?-selection. Sci Adv 9: eade5348

Chen L, Carico Z, Shih HY, Krangel MS (2015) A discrete chromatin loop in the mouse Tcra-Tcrd locus shapes the TCR8 and TCRα repertoires. Nat Immunol 16: 1085–1093

Cho WK, Jayanth N, English BP, Inoue T, Andrews JO, Conway W, Grimm JB, Spille J-H, Lavis LD, Lionnet T, Cisse II (2016) RNA polymerase II cluster dynamics predict mRNA output in living cells. eLife 5: e13617

Cho WK, Spille J-H, Hecht M, Lee C, Li C, Grube V, Cisse II (2018) Mediator and mRNA polymerase II clusters associate in transcription-dependent condensates. Science 361: 412–415

Cieslak A, Chanbonnier G, Tesio M, Mathieu E-L, Belhocine M, Touzart A, Smith C, Hypolite G, Andrieu GP, Martens JHA, Janssen-Megens E, Gut M, Gut I, Boissel N, Petit A, Puthier D, Macintyre E, Stunnenberg HG, Spicuglia S, Asnafi V (2020) Blueprint of human thymopoiesis reveals molecular mechanisms of stage-specific TCR enhancer activation. J Exp Med 217: e20192360

Cisse II, Izeddin I, Causse SZ, Boudarene L, Senecal A, Muresan L, Dugast-Darzacq C, Hajj B, Dahan M, Darzacq X (2013) Real-time dynamics of RNA polymerase II clustering in live human cells. Science 341: 664–667

da Costa Nunes JA, Noordermeer D (2023) TADs: Dynamic structures to create stably regulatory functions. Curr Opin Struct Biol 81: 102622

Dai R, Zhu Y, Li Z, Qin L, Liu N, Liao S, Hao B (2023) Three-way contact analysis characterizes the higher order organization of the Tcra locus. Nucleic Acids Res 51: 8987–9000

Dauphars DJ, Mihai A, Wang L, Zhuang Y, Krangel MS (2022) Trav15-dv6 family Tcrd rearrangements diversify the Tcra repertoire. J Exp Med 219: 20211581

de Wit E, Vos ESM, Holwerda SJB, Valdes-Quezada C, Verstegen MJAM, Teunissen H, Splinter E, Wijchers PJ, Krijger PHL, de Laat W (2015) CTCF binding polarity determines chromatin looping. Mol Cell 60: 676–684

del Blanco B, Angulo Ú, Krangel MS, Hernández-Munain C (2015) The Tcra enhancer is inactivated in α? T lymphocytes. Proc Natl Acad Sci U S A 112: 1744–1753

del Blanco B, García-Mariscal A, Wiest DL, Hernández-Munain C (2012) Tcra enhancer activation by inducible transcription factors downstream of pre-TCR signaling. J Immunol 188: 3278–3293

del Blanco B, Roberts JL, Zamarreño N, Balmelle-Devaux N, Hernández-Munain C (2009) Flexible and stereospecific interactions and composition within nucleoprotein complexes assembled on the TCR α gene enhancer. J Immunol 183: 1871–1883

Despang A, Schöpflin R, Franke M, Ali S, Jerkovic I, Paliou C, Chan W-L, Timmermann B, Wittler L, Vingron M, Mundlos S, Ibrahim DM (2019) Functional dissection of the Sox9-Kcnj2 locus identifies nonessential and instructive roles of TAD architecture. Nat Genet 51: 1263–1271

Dixon JR, Selvaraj S, Yue F, Kim A, Li Y, Shen Y, Hu M, Liu JS, Ren B (2012) Topological domains in mammalian genomes identified by analysis of chromatin interactions. Nature 485: 376–380

Dong Z, Wang H, Chen H, Jiang H, Yuan J, Yang Z, Wang W-J, Xu F, Guo X, Cao Y, Zhu Z, Geng C, Cheung WC, Kwok YK, Yang H, Leung TY, Morton CC, Cheung SW, Choy KW (2018) Identification of balanced chromosomal rearrangements previously unknown among participants in the 1000 Genomes Project: implications for interpretation of structural variation in genomes and the future of clinical cytogenetics. Genet Med 20: 697

Du M, Stizinger SH, Spille J-H, Cho W-K, Lee C, Hijad M, Quintana A, Cisse II (2024) Direct observation of a condensate effect on super-enhancer controlled gene bursting. Cell 187: 331–344

Espinola SM, Götz M, Bellec M, Messina O, Fiche J-B, Houbron C, Dejean M, Reim I, Casrdozo Gizzi AM, Lagha M, Nollman M (2021) Cis-regulatory chromatin loops arise before TADs and gene activation, and are independent of cell fate during early Drosophila development. Nat Genet 53: 477–486

Franke M, Ibrahim DM, Andrey G, Schwarzer W, Heinrich V, Schopflin R, Kraft K, Kempfer R, Jerkovic I, Chan W-L, Spielmann M, Timmermann B, Wittler L, Kurth I, Cambiaso P, Zuffardi O, Houge G, Lambie L, Brancati F, Pombo A, Vingron M, Spitz F, Mundlos S (2016) Formation of new chromatin domains determines pathogenicity of genomic duplications. Nature 538: 265–269

Freire-Pritchett P, Schoenfelder S, Várnai C, Wingett SW, Cairns J, Collier AJ, García-Vilchez R, Furlan-Magaril M, Osborne CS, Fraser P, Rugg-Gunn PJ, Spivakov M (2017) Global reorganization of cis-regulatory units upon lineage commitment of human embryonic stem cells. eLife 6: e21926

Fudenberg G, Imakaev M, Lu C, Goloborodko A, Abdennur N, Mirny LA (2016) Formation of chromosomal domains by loop extrusion. Cell reports 15: 2028–2049

Furlong EEM, Levine M (2018) Developmental enhancers and chromosome topology. Science 361: 1341–1345

Galupa R, Crocker J (2020) Enhancer-promoter communication: thinking outside the TAD. Trends Genet 36: 459–461

Galupa R, Nora EP, Worsley-Hunt R, Picard C, Gard C, van Bemmel JG, Servant N, Zhan Y, El Marjou F, Johanneau C, Diabangouaya P, Le Saux A, Lameiras S, Pipoli da Fonseca J, Loos F, Gribnau J, Baulande S, Ohler U, Giorgetti L, Heard E (2020) A conserved noncoding locus regulates random monoallelic Xist expression across a topological boundary. Mol Cell 77: 352–367

Galupa R, Picard C, Servant N, Nora EP, Zhan Y, van Bemmel JG, El Marjou F, Johanneau C, Borensztein M, Ancelin K, Giorgetti L, Heard E (2022) Inversion of a topological domain leads to restricted changes in its gene expression and affect interdomain communication. Development 149: dev200568

Ghavi-Helm Y, Jankowski A, Meiers S, Viales RR, Korbel JO, Furlong EEM (2019) Highly rearranged chromosomes reveal uncoupling between genome topology and gene expression. Nat Genet 51: 1272–1282

Ghavi-Helm Y, Klein FA, Padozdi T, Ciglar L, Noordermeer D, Huber W, Furlong EEM (2014) Enhancer loops appear stable during development and are associated with paused polymerase. Nature 512: 96–100

Guo Y, Xu Q, Canzio D, Shou J, Li J, Gorkin DU, Jung L, Wu H, Zhai Y, Tang Y, Lu Y, Wu Y, Jia Z, Li W, Zhang MQ, Ren B, Krainer AR, Maniatis T, Wu Q (2015) CRISPR inversion of CTCF sites alters genome topology and enhancer/promoter function. Cell 162: 900–910

Hagège H, Klous P, Braem C, Splinter E, Dekker J, Cathala G, de Laat W, Forné T (2007) Quantitative analysis of chromosome conformation capture assays (3C-qPCR). Nat Protoc 2: 1722–1733

Hernández-Munain C, Roberts JL, Krangel MS (1998) Cooperation among multiple transcription factors for access to minimal T-cell receptor α-enhancer chromatin in vivo. Mol Cell Biol 18: 3223–3233

Hernández-Munain C, Sleckman BP, Krangel MS (1999) A developmental switch from TCR 8 enhancer to TCR α enhancer function during thymocyte maturation. Immunity 10: 723–733

Hong NA, Flannery M, Hsieh SN, Cado D, Pedersen R, Winoto A (2000) Mice lacking Dad1, the defender against apoptotic death-1, express abnormal N-linked glycoproteins and undergo increased embryonic apoptosis. Dev Biol 220: 76–84

Hsieh T-HS, Cattoglio C, Slobodyanyuk E, Hansen AS, Darzacq X, Tjian R (2022) Enhancer-promoter interactions and transcription are largely maintained upon acute loss of CTCF, cohesin, WAPL or YY1. Nat Genet 54: 1919–1932

Hung T-C, Kingsley DM, Boettiger AN (2024) Boundary stacking interactions enable cross-TAD enhancer-promoter communication during limb development. Nat Genet 56: 306–314

Hussain S, Sadouni N, van Essen D, Dao LTM, Ferré Q, Charbonnier G, Torres M, Gallardo F, Lecellier C-H, Sexton T, Saccani S, Spicuglia S (2023) Short tandem repeats are important contributors to silence elements in T cells. Nucleic Acids Res 51: 4845–4866

Iborra FJ, Pombo A, Jackson DA, Cook PR (1996) Active RNA polymerases are localied within discrete transcription “factories” in human nuclei. J Cell Sci 109: 1427–1436

Ing-Simmons E, Vaid R, Bing XY, Levine M, Mannervik M, Vaquerizas JM (2021) Independence of chromatin conformation and gene regulation during Drosophila dorsoventral patterning. Nat Genet 53: 487–499

Irving BA, Alt FW, Killeen N (1998) Thymocyte development in the absence of pre-TCR receptor extracellular immunoglobulin domains. Science 280: 905–908

Javierre BM, Burren OS, Wilder SP, Kreuzhuber R, Hill SM, Sewitz S, Cairns J, Wingett SW, Várnai C, Thiecke MJ, Burden F, Farrow S, Cutler AJ, Rehnström K, Downes K, Grassi L, Kostadima M, Freire-Pritchett P, Wang F, Consortium B, Stunnenberg HG, Todd JA, Zerbino DR, Otegle O, Ouwehand WH, Frontini M, Wallace C, Spivakov M, Fraser P (2016) Lineage-specific genome architecture links enhancers and non-coding disease variants to target gene promoters. Cell 167: 1369–1384

Jeppsson K, Sakata T, Nakato R, Milanova S, Shirahige K, Björkegren C (2022) Cohesin-dependent chromosome loop extrusion is limited by transcription and stalled replication forks. Sci Adv 8: eabn7063

Kelleger DJ, Gilmore R (1997) DAD1, the defender against apoptotic cell death, is a subunit of the mammalian oligosaccharyltransferase. Proc Natl Acad Sci U S A 94: 4994–4999

Kessler S, Minoux M, Joshi O, Zouari YB, Ducret S, Ross F, Vuakin N, Salvi A, Wolff J, Kohler H, Stadler MB, Rijli FM (2023) A multiple super-enhancer region establishes inter-TAD interactions and controls Hoxa function in cranial neural crest. Nat Commun 14: 3242

Kim Y, Shi Z, Zhang H, Finkelstein IJ, Yu H (2019) Human cohesin compacts DNA by loop extrusion. Science 366: 1345–1349

Krijger PHL, Geeven G, Bianchi V, Hilvering CRE, de Laat W (2019) 4C-seq from beginning to end: A detailed protocol for sample preparation and data analysis. Methods 170: 17–32

Laugsch M, Bartusel M, Rehimi R, Alirzayeva H, Karaolidou A, Crispatzu G, Zentis P, Nikolic M, Bleckwehl T, Kolovos P, van Ijcken WFJ, Saric T, Koehler K, Frommolt P, Lachlan K, Baptista J, Rada-Iglesias A (2019) Modeling the pathological long-range regulatory effects of human structural variation with patient-specific hiPSCs. Cell Stem Cell 24: 736–752

Lawo S, Bashkurov M, Mullin M, Ferreria MG, Kittler R, Habermann B, Tagliaferro A, Poser I, Hutchins JR, Hegemann B, Pinchev D, Buchholz F, Peters JM, Hyman AA, Gingras AC, Pelletier L (2009) HAUS, the 8-subunit human Augmin complex, regulates centrosome and spindle integrity. Curr Biol 19: 816–826

Li J, Dong A, Saydaminova K, Chang H, Wang G, Ochiai H, Yamamoto T, Pertsinidis A (2019) Single-molecule nanoscopy elucidates RNA polymerase II transcription at single genes in live cells. Cell 178: 491–506

Lupiañez DG, Kraft K, Heinrich V, Krawitz P, Brancati F, Klopocki E, Horn D, Kayserili H, Opitz JM, Laxova R, Santos-Simarro F, Gilbert-Dussadier B, Wittler L, Borschiwer M, Haas SA, Osterwalder M, Frankle M, Timmermann B, Hetch J, Spielmann M, Visel A, Mundlos S (2015) Disruption of topological chromatin domains cause pathogenic rewiring of gene-enhancer interactions. Cell 161: 1012–1025

Mallis RJ, Bai K, Arthanari H, Hussey RE, Handley M, Li Z, Chingozha L, Duke-Cohan JS, Lu H, Wang JH, Zhu C, Wagner G, Reinherz EL (2015) Pre-TCR ligand binding impacts thymocyte development before α?TCR expression. Proc Natl Acad Sci U S A 112: 8373–8378

Mazzocca M, Narducci DN, Grosse-Holz S, Matthias J, Hansen AS (2025) Chromatin dynamics are highly subdiffusive across seven orders of magnitude. BioRxiv 2025.05.10.653248

Mihai A, Roy S, Krangel MS, Zhuang Y (2023) E protein binding at the Tcra enhancer promotes Tcra repertoire diversity. Front Immunol 14: 1188738

Naik AK, Dauphars D, Corbett E, Simpson L, Schatz DG, Krangel MS (2024) RORγt up-regulates RAG expression in DP thymocytes to expand the Tcra repertoire. Sci Immunol 9: eadh5318

Narendra V, Rocha PP, An D, Raviram R, Skok JA, Mazzoni EO, Reinberg D (2015) CTCF establishes discrete functional chromatin domains at the hox clusters during differentiation. Science 347: 1017–1021

Nora EP, Goloborodko A, Valton A-L, Gibcus JH, Uebersohn A, Abdennur N, Dekker J, Mirny LA, Bruneau BG (2017) Targeted degradation of CTCF decouples local insulation of chromosome domains from genomic compartimentalization. Cell 169: 930–944

Nora EP, Lajoie BR, Schultz EG, Giorgetti L, Okamoto I, Servant N, Piolot T, van Berkum NL, Meisig J, Sedat J, Gribnau J, Barillot E, Blüthgen N, Dekker J, Heard E (2012) Spatial partitioning of the regulatory landscape of the X-inactivation centre. Nature 485: 381–385

Phillips-Cremins JE, Sauria MEG, Sanyal A, Gerasimova TI, Lajoie BR, Bell JSK, Ong C-T, Hookway TA, Guo C, Sun Y, Bland MJ, Wagstaff W, Dalton S, McDevitt TC, Sen R, Dekker J, Taylor J, Corcés VG (2013) Architectural protein subclasses shape 3D organization of genomes during lineage commitment. Cell 153: 1281–1295

Rao SS, Huntley MH, Durand NC, Stamenova EK, Bochkov ID, Robinson JT, Sanborn AL, Machol I, Omer AD, Lander ES, Aiden EL (2014) A 3D map of the human genome at kilobase resolution reveals principles of chromatin looping. Cell 159: 1665–1680

Rao SSP, Huang S-C, Glen St Hilarie B, Engreitz JM, Perez EM, Kieffer-Kwon KR, Sanborn AL, Johnstone SE, Bascom GD, Bochkov ID, Huang X, Shamin MS, Shin J, Turner D, Ye Z, Omer AD, Robinson JT, Schlick T, Bernstein BE, Casellas R, Lander ES, Aiden EL (2017) Cohesin loss eliminates all loop domains. Cell 171: 305–320

Rodríguez-Caparrós A, Álvarez-Santiago J, López-Castellanos L, Ruiz-Rodríguez C, Valle-Pastor MJ, López-Ros J, Angulo Ú, Andrés-León E, Suñé C, Hernández-Munain C (2022) Differently regulated gene-specific activity of enhancers located at the boundary of sub-topologically associated domains: TCRα enhancer. J Immunol 208: 910–928

Rodríguez-Caparrós A, Álvarez-Santiago J, Valle-Pastor MJ, Suñé C, López-Ros J, Hernández-Munain C (2020) Regulation of T-cell receptor gene expression by three-dimensional locus conformation and enhancer function. Int J Mol Sci 21: 8478

Rodríguez-Caparrós A, García V, Casal Á, López-Ros J, García-Mariscal A, Tani-ichi S, Ikuta K, Hernández-Munain C (2019) Notch signaling controls transcription via the recruitment of RUNX1 and MYB to enhancers during thymocyte development. J Immunol 202: 2460–2472

Rodríguez-Carballo E, López-Delisle L, Zhan Y, Fabre PJ, Beccari L, El-Idrissi I, Huynh THN, Ozadam H, Dekker J, Duboule D (2017) The HoxD cluster is a dynamic and resilient TAD boundary controlling the segregation of antagonistic regulatory landscapes. Genes Dev 31: 2264–2281

Rouco R, Bompadre O, Rauseo A, Fazio O, Peraldi R, Thorel F, Andrey G (2021) Cell-specific alterations in Pitx1 regulatory landscape activation caused by loss of a single enhancer. Nat Commun 12: 7235

Sanborn AL, Rao SS, Huang SC, Durand NC, Huntley MH, Jewett AI, Bochkov ID, Chinappan D, Cutkosky A, Li J, Geeting KP, Gnirke A, Melnikov A, McKenna D, Stamenova EK, Lander ES, Aiden EL (2015) Chromatin extrusion explains key features of loop and domain formation in wild-type and engineered genomes. Proc Natl Acad Sci U S A 112: E6456–6465

Sanyal A, Lajoie BR, Jain G, Dekker J (2012) The long-range interaction landscape of gene promoters. Nature 489: 109–113

Schönberg PY, Muñoz-Ovalle Á, Paszkowski-Rogacz M, Crespo E, Sürün D, Feldmann A, Buchholz F (2025) A pooled CRISPR screen identifies the Tα2 enhancer element as a driver of TRA expression in a subset of mature T lymphocytes. Front Immunol 16: 1536003

Schwarzer W, Abdennur N, Goloborodko A, Pekowska A, Fudenberg G, Loe-Mie Y, Fonseca NA, Huber W, Haering CH, Mirny LA, Spitz F (2017) Two independent modes of chromatin organization revealed by cohesin removal. Nature 551: 51–56

Seitan VC, Faure AJ, Zhan Y, McCord RP, Lajoie BR, Ing-Simmons E, Lenhard B, Giorgetti L, Heard E, Fisher AG, Flicek P, Dekker J, Merkenschlager M (2013) Cohesin-based chromatin interactions enable regulated gene expression within preexisting architectural comartments. Genome Res 23: 2066–2077

Shen Y, Yue F, McCleary DF, Ye Z, Edsall L, Kuan S, Wagner U, Dixon J, Lee L, Lobanenkov VV, Ren B (2012) A map of the cis-regulatory sequences in the mouse genome. Nature 488: 116–120

Shih HY, Verma-Gaur J, Torkamani A, Feeney AJ, Galjart N, Krangel MS (2012) Tcra gene recombination is supported by a Tcra enhancer-and CTCF-dependent chromatin hub. Proc Natl Acad Sci U S A 109: 3493–3502

Shinkai Y, Alt FW (1994) CD3β-mediated signals rescue the development of CD4+CD8+ thymocytes in RAG-2-/- mice in the absence of TCR? chain expression. Int Immunol 6: 995–1001

Shinkai Y, Rathbun G, Lam P, Oltz EM, Stewart V, Mendelsohn M, Charron J, Datta M, Young F, Stall AM, Alt FW (1992) RAG-2-deficient mice lack mature lymphocytes owing to inability to initiate V(D)J recombination. Cell 68: 855–867

Sleckman BP, Bardon CG, Ferrini R, Davidson L, Alt FW (1997) Function of the TCR α enhancer in α? and Ψ8 T cells. Immunity 7: 505–515

Smith EM, Lajoie BR, Jain G, Dekker J (2016) Invariant TAD boundaries constrain cell-type-specific looping interactions between promoters and distal elements around the CFTR locus. AJHG 98: 185–201

Soshsnikova N, Montavon T, Leleu M, Galjart N, Duboule D (2017) Functional analysis of CTCF during mammalian limb development. Dev Cell 19: 819–830

Spicuglia S, Payet D, Tripathi RK, Rameil P, Verthuy C, Imbert J, Ferrier P, Hempel WM (2000) TCRα enhancer activation occurs via a conformational change of a pre-assembled nucleo-protein complex. EMBO J 20: 42–53

Sun F, Chronis C, Kronenberg M, Chen XF, Su T, Lay FD, Plath K, Kurdistani SK, Carey MF (2019) Promoter-enhancer communication occur primarily within insulated neighborhoods. Mol Cell 73: 250–263

Symmons O, Uslu VV, Tsujimura T, Ruf S, Nassari S, Schwarzer W, Ettwiller L, Spitz F (2014) Functional and topological characteristics of mammalian regulatory domains. Genome Res 24: 390–400

van Arensbergen J, van Steensel B, Bussemaker HJ (2014) In search of the determinants of enhancer-promoter interaction specificity. Trends Cell Biol 24: 695–702

Vanhille L, Griffon A, Maqbool MA, Zacarías-Cabeza J, Dao LT, Fernandez N, Ballester B, Andrau JC, Spicuglia S (2015) High-through and quantitative assesment of enhancer activity in mammals by CapStarr-seq. Nat Commun 6: 6905

Wang H, Li B, Zuo L, Wang B, Yan Y, Tian K, Zhou R, Wang C, Chen X, Jiang Y, Zheng H, Qin F (2022) The transcriptional coactivator RUVBL2 regulates Pol II clustering with diverse transcription factors. Nat Commun 13: 5703

Wutz G, Várnai C, Nagasaka K, Cisneros DA, Stocsits RR, Tang W, Schoenfelder S, Jessberger G, Muhar M, Hossain MJ, Walther N, Koch B, Kueblbeck M, Ellenberg J, Zuber J, Fraser P, Peters JM (2017) Topologically associating domains and chromatin loops depend on cohesin and are regulated by CTCF, WAPL, and PDS5 proteins. EMBO J 36: 3573–3599

Yang JH, Hansen AS (2024) Enhancer selectively in space and time: From enhancer-promoter interactions to promoter activation. Nat Rev Mol Cell Biol 25: 574–591

Zhan Y, Mariani L, Barozzi I, Schulz EG, Blütgen N, Stadler M, Tiana G, Giorgetti L (2017) Reciprocal insulation analysis of Hi-C data shows that TADs represent a functionally but not structurally privileged scale of the hierarchical folding of chromosomes. Genome Res 27: 479–490

Zhao H, Li Z, Zhu Y, Bian S, Zhang Y, Qin L, Naik AK, He J, Zhang Z, Krangel MS, Hao B (2020) A role for CTCF binding site at enhancer Eα in the dynamic chromatin organization of the Tcra-Tcrd locus. Nucleic Acids Res 48: 9621–9636

Zuin J, Dixon JR, van der Reijden MIJ, Ye Z, Kolovos P, Brouwer RWW, van de Corput MPC, van de Werken HJG, Knoch TA, van Ijcken WFJ, Grosveld F, Ren B, Wendt KS (2014) Cohesin and CTCF differentially affect chromatin architecture and gene expression in human cells. Proc Natl Acad Sci U S A 111: 996–1001

